# Introduction of *loxP* sites by electroporation in the mouse genome; a simple approach for conditional allele generation in complex targeting loci

**DOI:** 10.1101/2021.12.14.471503

**Authors:** Guillaume Bernas, Mariette Ouellet, Andréa Barrios, Hélène Jamann, Catherine Larochelle, Émile Lévy, Jean-François Schmouth

**Affiliations:** Centre de recherche du CHUM, Université de Montréal, Montréal, Canada; Département de Neurosciences, Université de Montréal, Montréal, Canada; Centre de recherche du CHU Ste-Justine, Université de Montréal, Montréal, Canada; Département de Pharmacologie et physiologie; Département de Nutrition

**Keywords:** CRISPR, Electroporation, conditional, *loxP*, zygotes

## Abstract

**Background:** The discovery of the CRISPR-Cas9 system and its applicability in mammalian embryos has revolutionized the way we generate genetically engineered animal models. To date, models harbouring conditional alleles (i.e.: two *loxP* sites flanking an exon or a critical DNA sequence of interest) remain the most challenging to generate as they require simultaneous cleavage of the genome using two guides in order to properly integrate the repair template. In the current manuscript, we describe a modification of the sequential electroporation procedure described by Horii *et al* (2017). We demonstrate production of conditional allele mouse models for eight different genes via one of two alternative strategies: either by consecutive sequential electroporation (strategy A) or non-consecutive sequential electroporation (strategy B).

**Results:** By using strategy A, we demonstrated successful generation of conditional allele models for three different genes (*Icam1, Lox*, and *Sar1b*), with targeting efficiencies varying between 5 to 13%. By using strategy B, we generated five conditional allele models (*Loxl1, Pard6a, Pard6g, Clcf1*, and *Mapkapk5*), with targeting efficiencies varying between 3 to 25%.

**Conclusion:** Our modified electroporation-based approach, involving one of the two alternative strategies, allowed the production of conditional allele models for eight different genes via two different possible paths. This reproducible method will serve as another reliable approach in addition to other well-established methodologies in the literature for conditional allele mouse model generation.

## Background

The discovery of the CRISPR-Cas9 system and its applicability in mammalian embryos has revolutionized the way we generate genetically engineered animal models. The generation of models relies on the delivery of CRISPR-Cas9 components in embryos that results in induction of double strand DNA breaks at predefined-specific sites in the genome [1]. Intrinsic mammalian DNA repair mechanisms are then used to either ablate exons or insert random mutations such as insertions and deletions (indels), via the non-homologous end joining (NHEJ) pathway, or introduce specific DNA repair templates, harboring point mutations, targeted reporters or conditional alleles via the homology dependent repair (HDR) pathway [2]. In rodent production, the delivery process relies on two main approaches, either by microinjection (into the pronucleus or nucleus) or by electroporation. The microinjection procedure consists of microinjecting CRISPR-Cas9 components either into the cytoplasm or directly into one of the two pronuclei of 1-cell embryos or one or both nuclei of 2-cell embryos [3, 4]. This approach has the advantage of being highly versatile as it can be applied to generate any type of model. However, it relies on the use of expensive microscopy setups and highly trained personnel. For these reasons, microinjection is mainly used by centralized cores and is usually offered as a service. On the other hand, electroporation consists of using an electrical current to open up pores in embryo membranes to allow the entry of CRISPR-Cas9 components [5, 6]. This approach has the advantage of being less technically challenging and does not require expensive microscopy setups, making it more appealing to individual laboratories. The drawback of this approach is that its widespread applicability is restricted to classical knockout and point mutation alleles as the size of DNA constructs that can be incorporated through the opened pore is limited.

To date, models harbouring conditional alleles (i.e.: two *loxP* sites flanking an exon or a critical DNA sequence of interest) remain the most challenging to generate as they require the simultaneous cleavage of two guides in order to properly integrate the repair template. One published approach to generate these types of alleles is the *Easi*-CRISPR method, which employs a long single stranded DNA (lssDNA) repair template that is injected concurrently with the CRISPR-Cas9 components in mouse pronucleus zygotes [7, 8]. This allows the incorporation of two *loxP* sites surrounding the desired sequence, at a specific locus in the genome. The microinjection procedure is performed on 1-day fertilized embryos that are then implanted in pseudopregnant females, and the resulting pups are characterized for proper *loxP* sites integration. A similar approach, named CRISPR with lssDNA inducing conditional knockout allele (CLICK), has been reported and uses lssDNA repair templates to generate conditional allele by electroporation [9]. This method, although successful in generating conditional alleles with repair templates of up to 1.4 kb, has the drawback of requiring a large amount of lssDNA. This makes it less appealing to centralized cores since these constructs are generally provided by commercial vendors, are expensive, and in limited supply. As a transgenic core, we have successfully used the *Easi*-CRISPR method. However, we have also experienced limitations with this approach. For example, synthesis of the required lssDNA construct by commercial vendors is usually restricted to less than 2 kb, rendering this approach unsuitable for projects targeting multiple or large exons. In addition, in some instances, sequence complexity hinders lssDNA synthesis, further restricting the flexibility of this approach. Finally, the high price of commercially-produced lssDNA constructs (provided in limited amount) makes this approach less appealing for projects where budget is a limiting factor.

An alternative method has been described where electroporation is used to incorporate the CRISPR-Cas9 components and two short single strand oligonucleotides (ssODNs) into mouse zygotes in order to integrate the *loxP* sites at a specific locus. This is achieved by two rounds of electroporation, one at the one cell stage (1-day fertilized embryo) and the second round performed 24 hours later, at the two cells stage. The electroporated embryos are then implanted into pseudopregrant females and the resulting pups are characterized for proper incorporation of the two *loxP* sites [10].

In the current manuscript, we have adapted this method to both a consecutive sequential (strategy A) and non-consecutive sequential (strategy B) electroporation method to generate novel mouse models with conditional alleles. We demonstrate that the use of two ssODNs in consecutive and non-consecutive sequential electroporation to introduce *loxP* sites is a reliable and flexible method that should be considered as an alternative approach to other methods currently used. Moreover, we have successfully applied this modified approach to the generation of animal models with *loxP* sites several kbs apart or with sequences that were too complex for commercial synthesis; two limitations that would have otherwise made these projects impossible using a lssDNA as a repair template. Moreover, this method provides budgetary flexibility when considering the guide and repair template choices as well as highlights two different path (strategy A and strategy B) leading to the successful generation of the desired conditional knockout mouse model.

## Results

### Applying consecutive or non-consecutive sequential electroporations to generate novel conditional alleles: a roadmap

As a transgenic core, we have faced the challenge of receiving a large number of requests for novel conditional allele models in a short period of time. To date, the *Easi-*CRISPR method, employing a combination of two guide RNAs complexed with the Cas9 protein along with a lssDNA repair template, seems to be the most widely adapted method for generating conditional alleles. The method has been proven to be robust and reliable for generating conditional alleles for most genes and was successfully applied for two previously generated models in our laboratory (data not shown) [8]. However, during the course of our work, we realized some limitations of this approach that precluded its applicability to all loci (details see table 1). Specifically, the *Easi*-CRISPR method was unusable for four out of eight projects (details see table 1). The projects involving *Icam1* and *Clcf1* were incompatible with the *Easi*-CRISPR approach due to the distance between both *loxP* sites, requiring a targeting construct greater than 2 kb. Moreover, *Easi*-CRISPR could not be successfully applied to the projects involving *Lox* and *Pard6g* due to the high sequence complexity surrounding the targeting region. To circumvent these limitations, we used a modified version of the electroporation conditions reported by Troder *et al*. along with the sequential electroporation method reported by Horii et *al* (details see methods section) [10, 11]. The rationale that we have used for each project is summarized in Figure 1. In short, we employed a strategy to generate conditional alleles according to two possible scenarios; 1) by consecutive sequential electroporation (strategy A); or 2) by non-consecutive sequential electroporation (strategy B). The first attempt for each project was via consecutive sequential electroporation (Strategy A, blue rectangle Box, figure 1). We rationalized that this approach was the shortest path to success if it worked. If it failed, we investigated whether or not any pups resulting from the initial consecutive sequential electroporation session could be usable for the non-consecutive sequential electroporation approach (Strategy B, grey rectangle Box, figure 1).

**Table 1:**
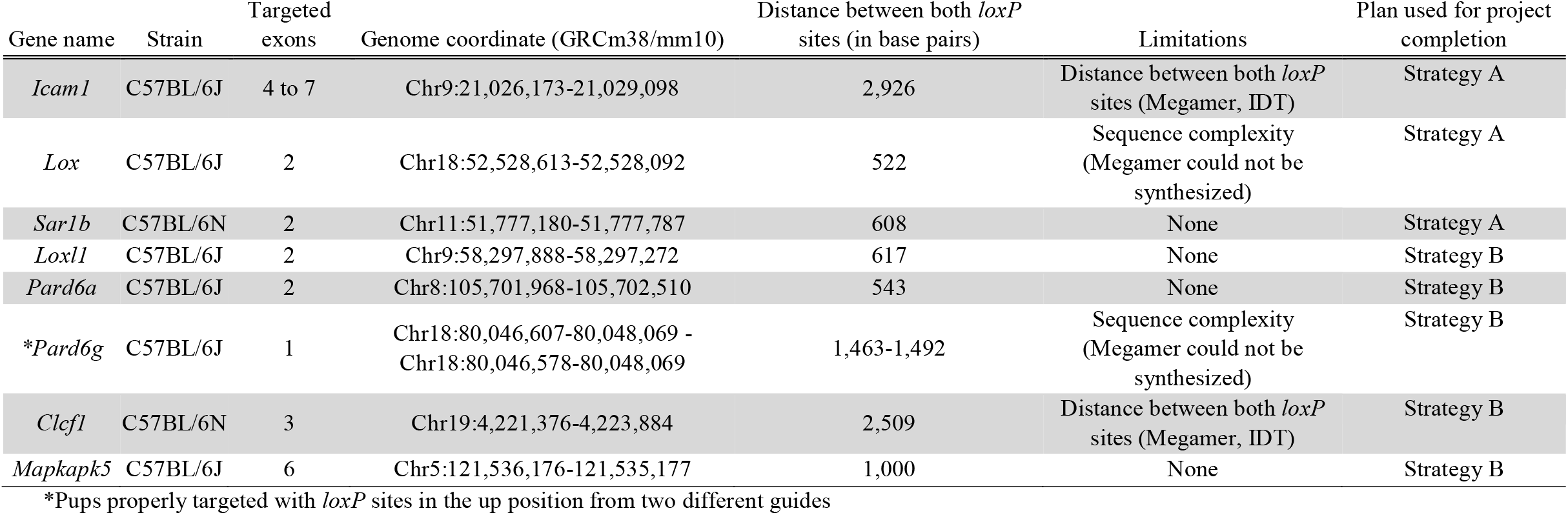
Details of the conditional allele targeting projects. All eight different conditional allele projects mentioned in this manuscript are highlighted in the current table. Column 1 identify the targeted genes (common name identifier), column 2 highlights the strain background used, column 3 highlights the exon(s) targeted for each project, column 4 highlights the genome coordinates according to the UCSC-GRCm38/mm10 mouse genome assembly, column 5 highlights the distance between both *loxP* sites, column 6 highlights the limitations encountered for each project, and column 7 highlights the strategy used for project completion.

**Figure 1:**
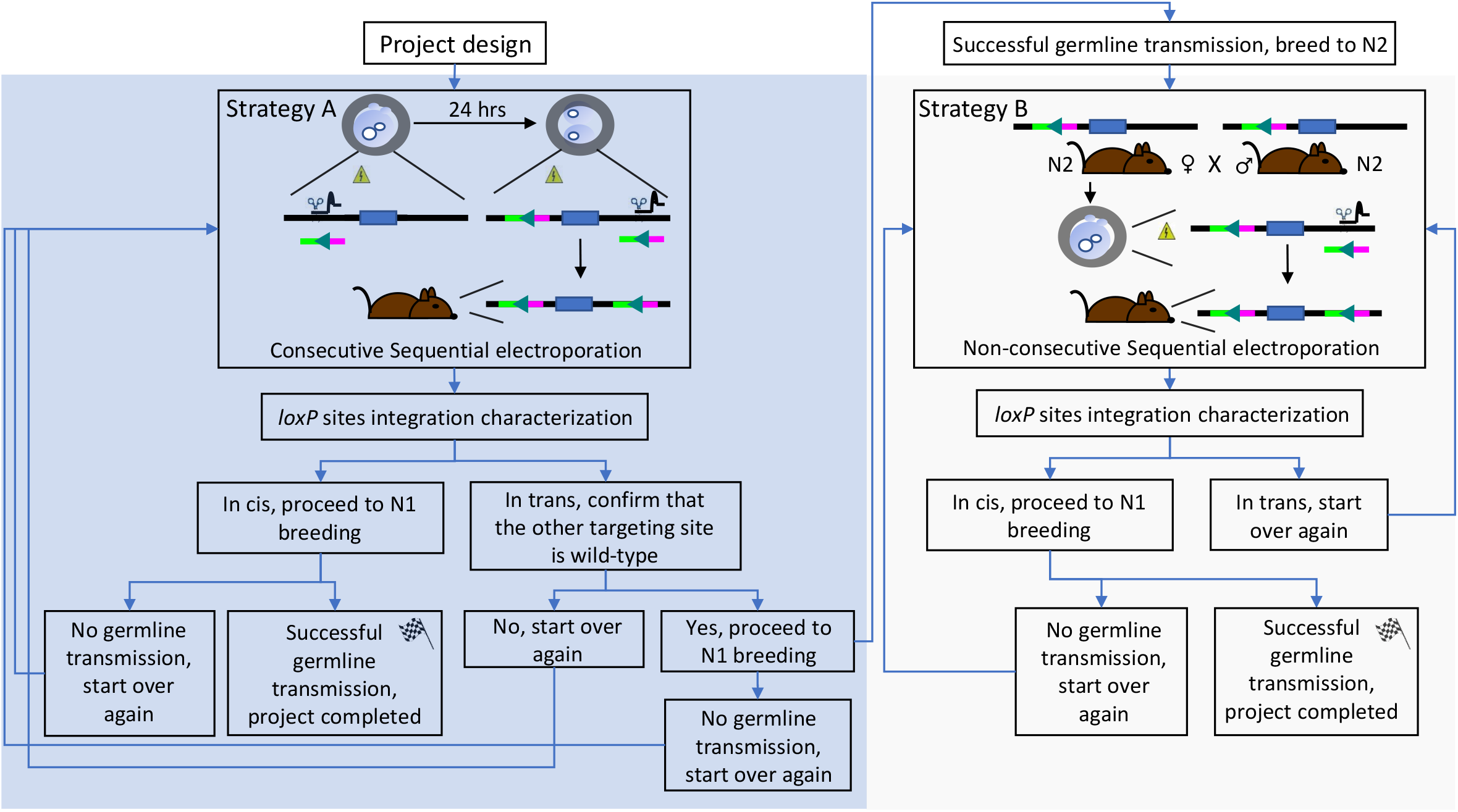
Decision three highlighting the different options leading to successful conditional allele generation. A decision three representing the different options leading to successful conditional allele generation is represented. The project success is based on two different scenarios depending on the initial electroporation outcomes, either by consecutive sequential electroporation (Strategy A) or non-consecutive sequential electroporation (Strategy B).

This modified procedure was extended to the four remaining projects highlighted in table 1. These included two projects where the *Easi*-CRISPR method failed to produce animals containing the desired alleles (*Sar1b* and *Loxl1*, table 1 and additional file 1). In this case, three positive animals were obtained that contained partial construct integrations at the targeted site (one for *Sar1b* and two for *Loxl1*, additional file 1) and five animals were obtained that contained random construct integrations (three for *Sar1b* and two for *Loxl1*, additional file 1). Random and partial integration screening strategies for *Sar1b* and *Loxl1* are detailed in additional files 2-3. The *Sar1b* partially integrated construct contained a properly targeted *loxP* site on one side of the desired exon and a 14 base pairs deletion on the opposite side. The *Loxl1* partial integration consisted of a sequence inversion in the 3’homology arm of the repair template along with a 40 base pairs deletion for one animal and a properly targeted *loxP* site on one side of the desired exon with no indels on the opposite side. This latter observation suggested a difference in guide cleaving efficiency for this project. Interestingly, these two phenomena of random and partial integrations have been previously reported in the literature for projects using lssDNA [7, 12]. We extended our consecutive and non-consecutive electroporation strategies to the two remaining projects that had no limitation for using lssDNA as a repair template (*Pard6a* and *Mapkapk5*, table 1). For these two projects, the consideration of cost and synthesis turnover time for the generation of a lssDNA construct weighed against the possibility of obtaining a partial integration model, prompted us to instead invest in short ssODNs. Our rationale also took into consideration the fact that in most cases, a single properly targeted *loxP* site animal with wild-type sequence on the opposite site could be obtained leading us to a strategy B alternative (non-consecutive sequential electroporation). Essentially, in this case, we reasoned that one properly targeted *loxP* site was better than none.

### Applying consecutive or non-consecutive sequential electroporation strategies to generate novel conditional allele: project design

For each project, the design relied on the selection and use of two annealed RNA guides, referred here as pgRNA (crRNA-tracrRNA formulation) and symmetric short single strand oligonucleotides (ssODNs) as repair templates that contained 60 base pairs homology arms on each side, and a *loxP* sequence in between (repair template details, see table 2). Sequence length between both homology arms varied depending on whether a single *loxP* site (34 base pairs) or an associated adjacent *EcoRI* or *NheI* restriction sites (40 base pairs) was incorporated along the *loxP* sequence. The repair templates were designed to correspond to the targeting strand, complementary to the Cas9 selected guide and its associated PAM site sequence, with an exception for the *Loxl1* project, where repair templates of both orientations in the Dn position (3’ of the targeted exon) were used to complete the project (table 2).

**Table 2:**
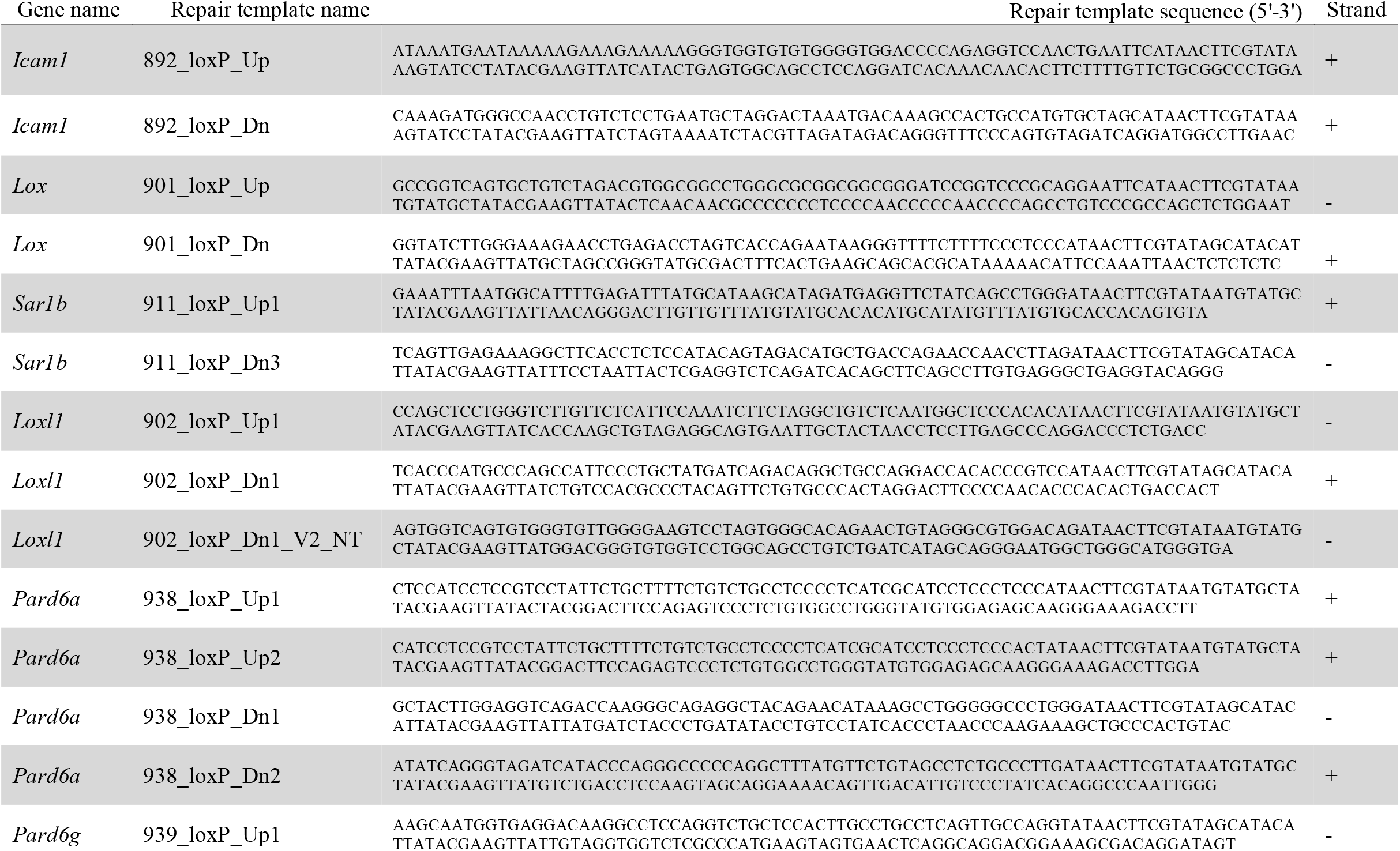
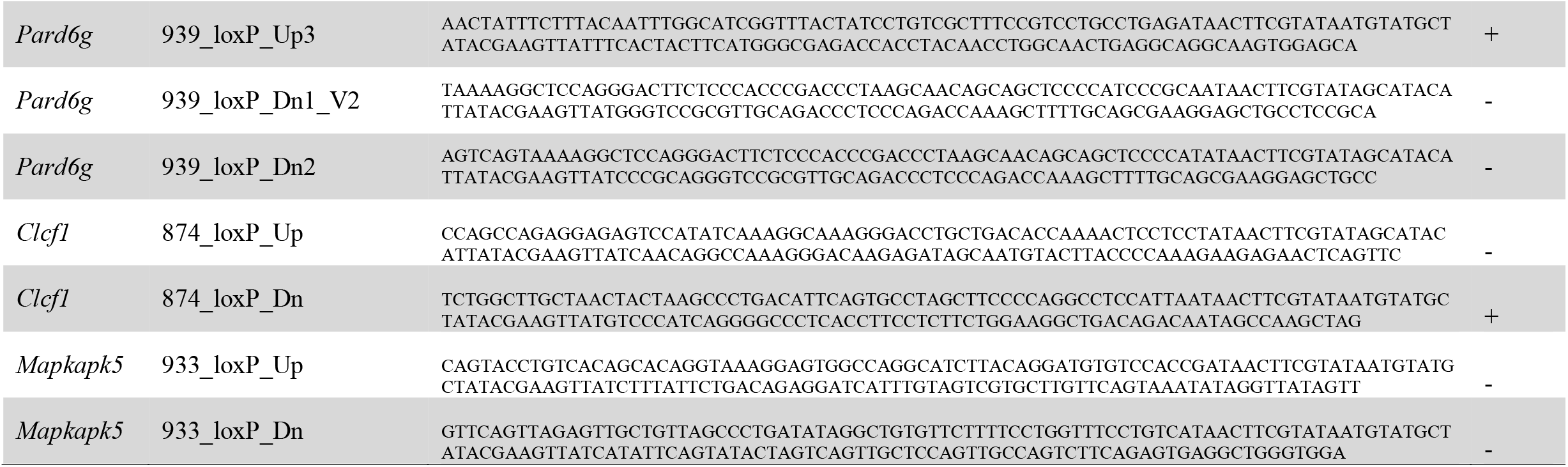
Repair template details for all eight projects. The repair templates designed for each project are highlighted in the current table. Column 1 highlights the gene names, column 2 highlights the repair template names, column 3 highlights the repair template sequences and column 4 highlights the corresponding strand orientation in the genome (according to the UCSC-GRCm38/mm10 mouse assembly).

Literature review and gene structure analyses were performed for each individual project to select exons that were predicted to have the most detrimental effect on the protein product when deleted. *In silico* guide cutting surveys were performed for each candidate exon using three different softwares (CRISPOR (http://crispor.tefor.net/crispor.py), CHOPCHOP (https://chopchop.cbu.uib.no/) and Breaking-Cas (https://bioinfogp.cnb.csic.es/tools/breakingcas/)) on selected genomic DNA regions as described in the methods section [13-15]. A total of three guide cutting pairs were selected for each individual project. crRNAs corresponding to the top-ranking guide pair, cutting on each side of the candidate(s) exon(s), were ordered from IDT along with the two corresponding ssODN repair templates. Complete lists of the different crRNA and corresponding repair templates are highlighted in table 2 and 3. In some instances, an additional crRNA pair was ordered and used in the initial sequential electroporation procedure (*Pard6a* and *Pard6g*, table 2). Consecutive sequential electroporation sessions were performed for each selected guide pairs as described in the methods section. Briefly, RNP complexes formed by the association of one of the two selected pgRNA with the purified Cas9 protein were electroporated in 1-cell stage embryos along with the corresponding repair template (Strategy A, figure 1). Electroporated embryos were recovered and left to develop to the 2-cells stage overnight at 37°C under 5% CO_2_. 2-cells stage embryos were electroporated with the second RNP complex along with the corresponding repair template before being implanted in pseudopregnant females (0.5 dpc) (Strategy A, figure 1).

**Table 3:**
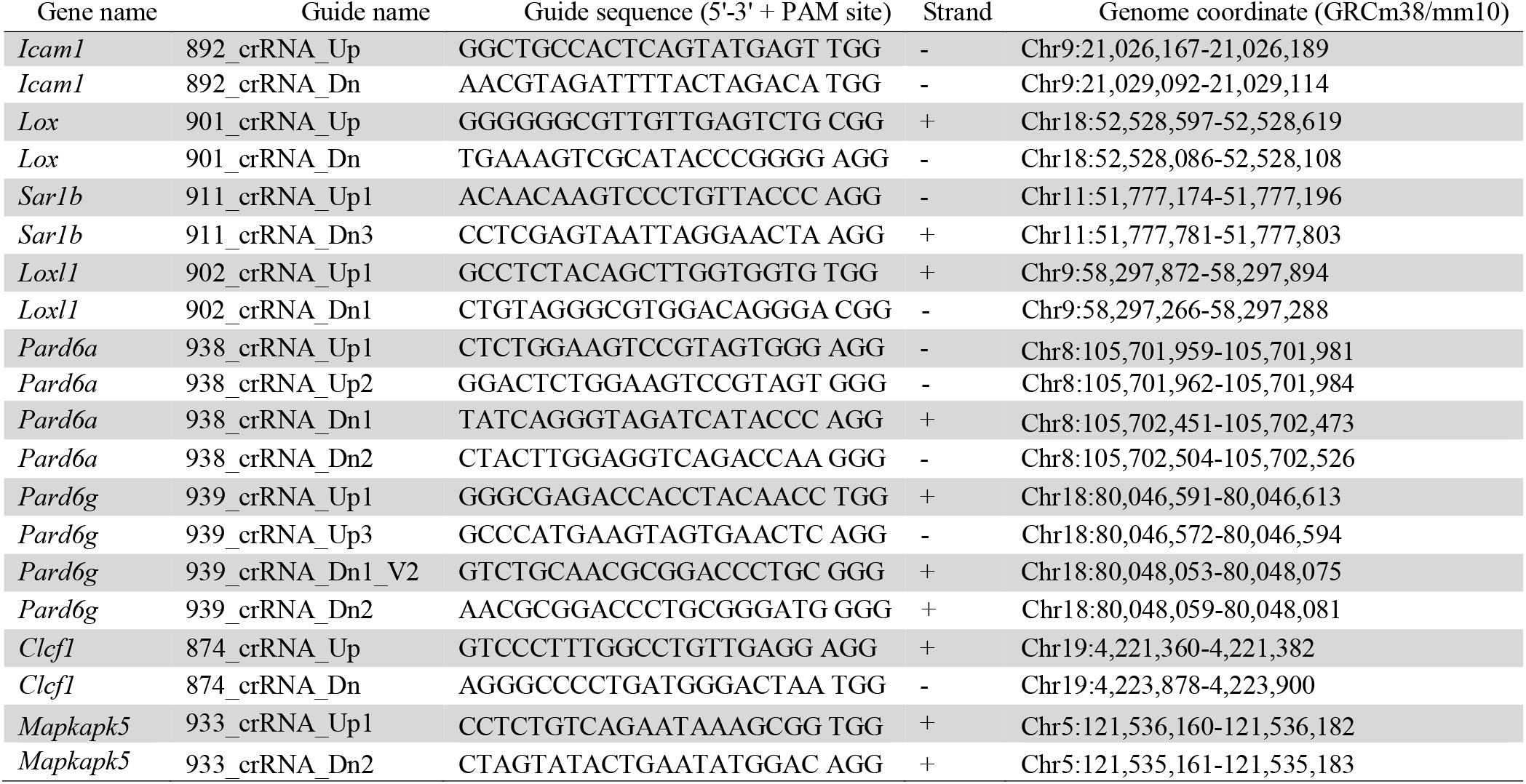
Guide sequence details for all eight projects. The guide sequences selected for each project are highlighted in the current table. Column 1 highlights the gene names, column 2 highlights the guide names, column 3 highlights the guide sequences, column 4 highlights the corresponding strand orientations in the genome and column 5 highlights the corresponding genome coordinates according to the UCSC-GRCm38/mm10 mouse assembly.

### Applying consecutive or non-consecutive sequential electroporations to generate novel conditional allele: properly targeted pups characterization

The resulting pups were characterized using a genotyping approach previously described in the literature with primer series exemplified in blue in figure 2A [8]. Briefly, six primers were routinely designed for each project. These comprised two pairs, mapping outside the ssODN homology arms used to insert the *loxP* site either in the Up (5’ of the targeted exon(s)) or Dn (3’ of the targeted exon(s)) positions (Figure 2A, primers 1-3, Up; primers 4-6, Dn). Two additional primers were designed with overlaps between the genomic DNA sequence adjacent to the *loxP* insertion site (20 base pairs) and a portion (15 base pairs) of the *loxP* site itself (Figure 2A, primers 2 and 5). These last primers were designed to be used as a pair with one primer pointing in the forward and the other in the reverse orientation. A complete primer list for each project is found in table 4.

**Table 4:**
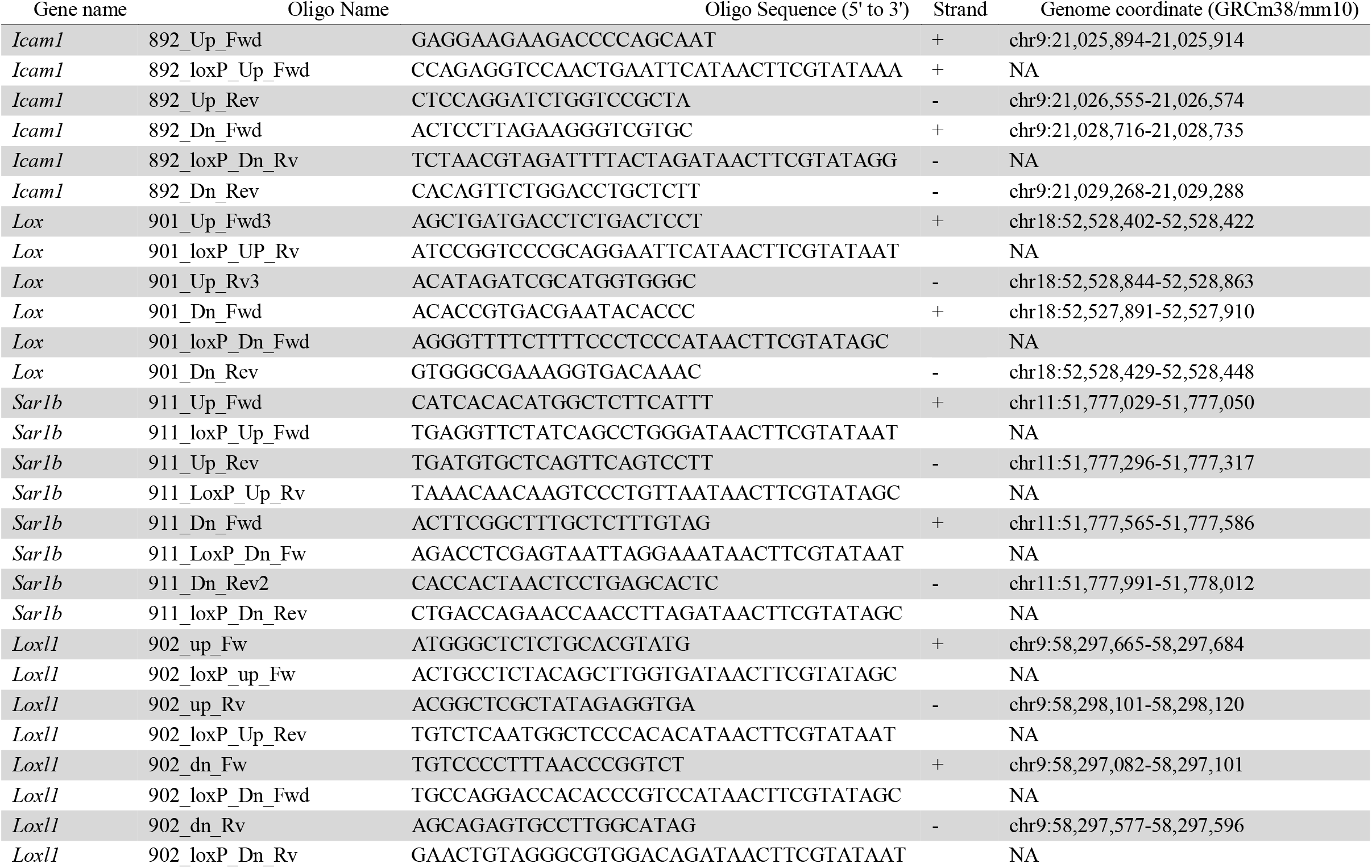
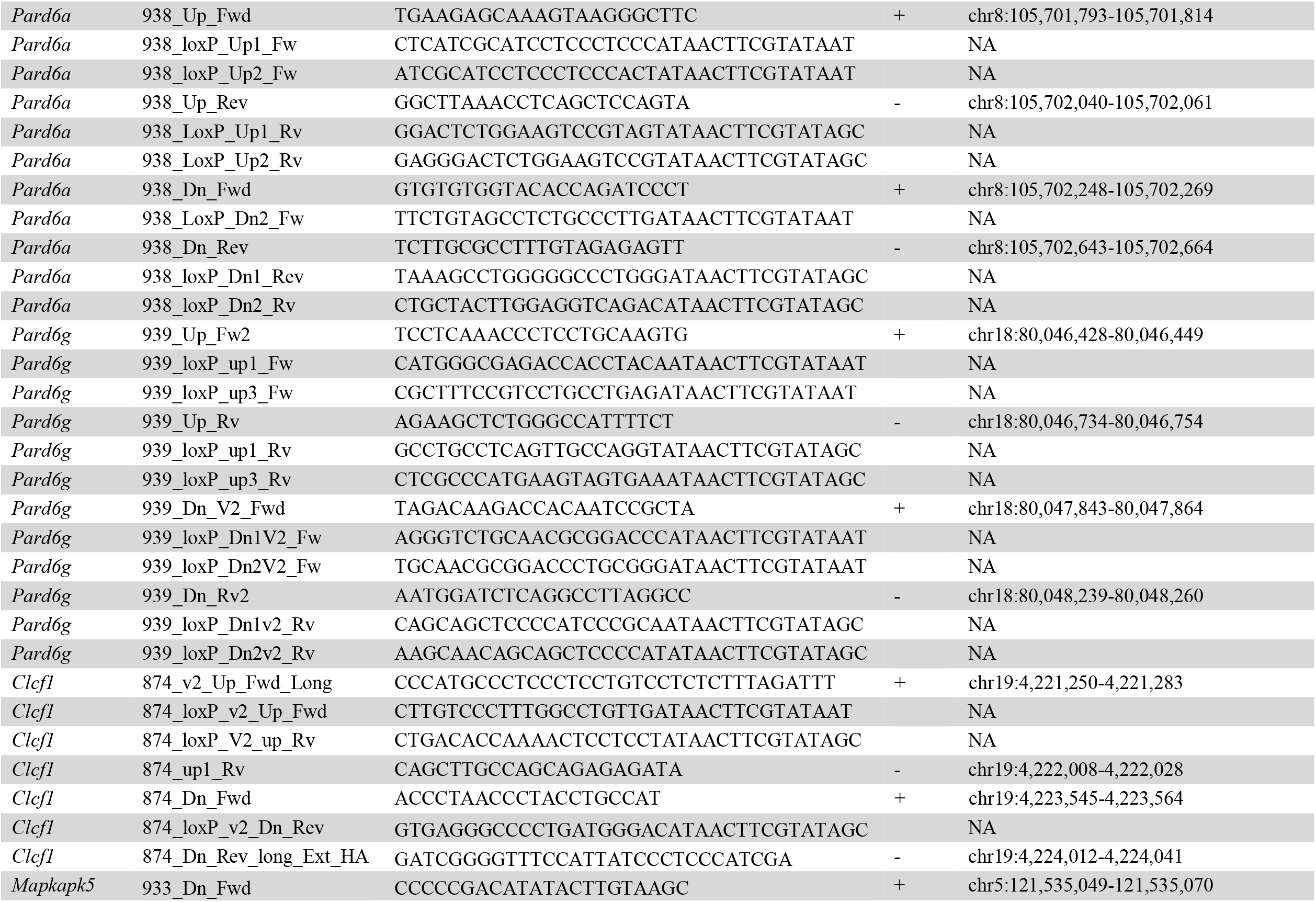
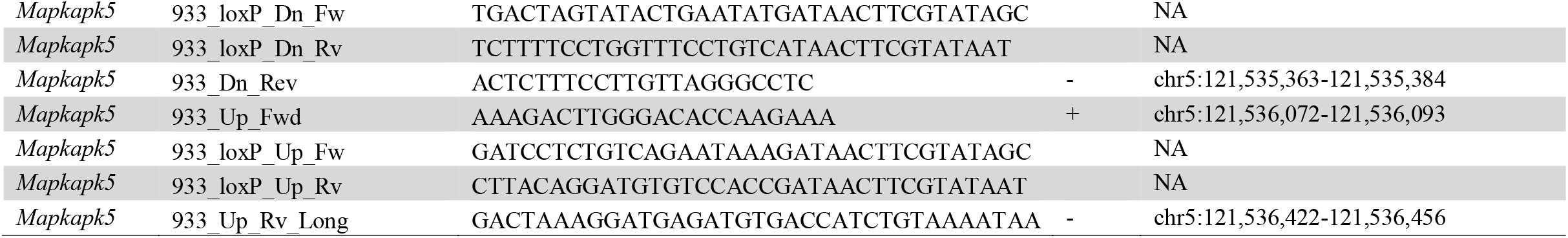
PCR oligo details for all eight projects. The PCR oligos for each project are highlighted in the current table. Column 1 highlights the gene name, column 2 highlights the PCR oligo names, column 3 highlights the oligo sequences, column 4 highlights the corresponding strand orientation in the genome and column 5 highlights the corresponding genome coordinates according to the UCSC-GRCm38/mm10 mouse assembly.

**Figure 2:**
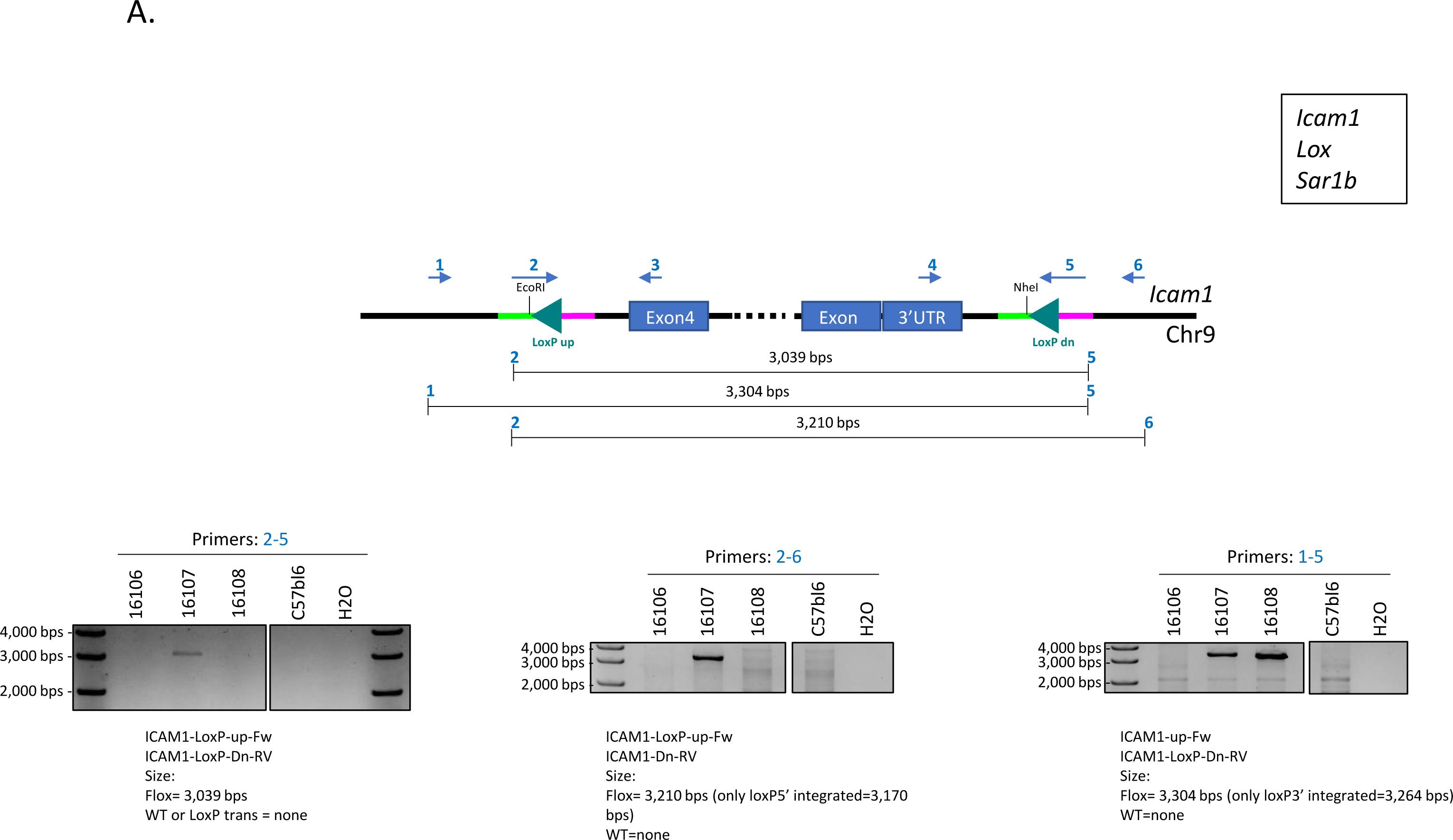
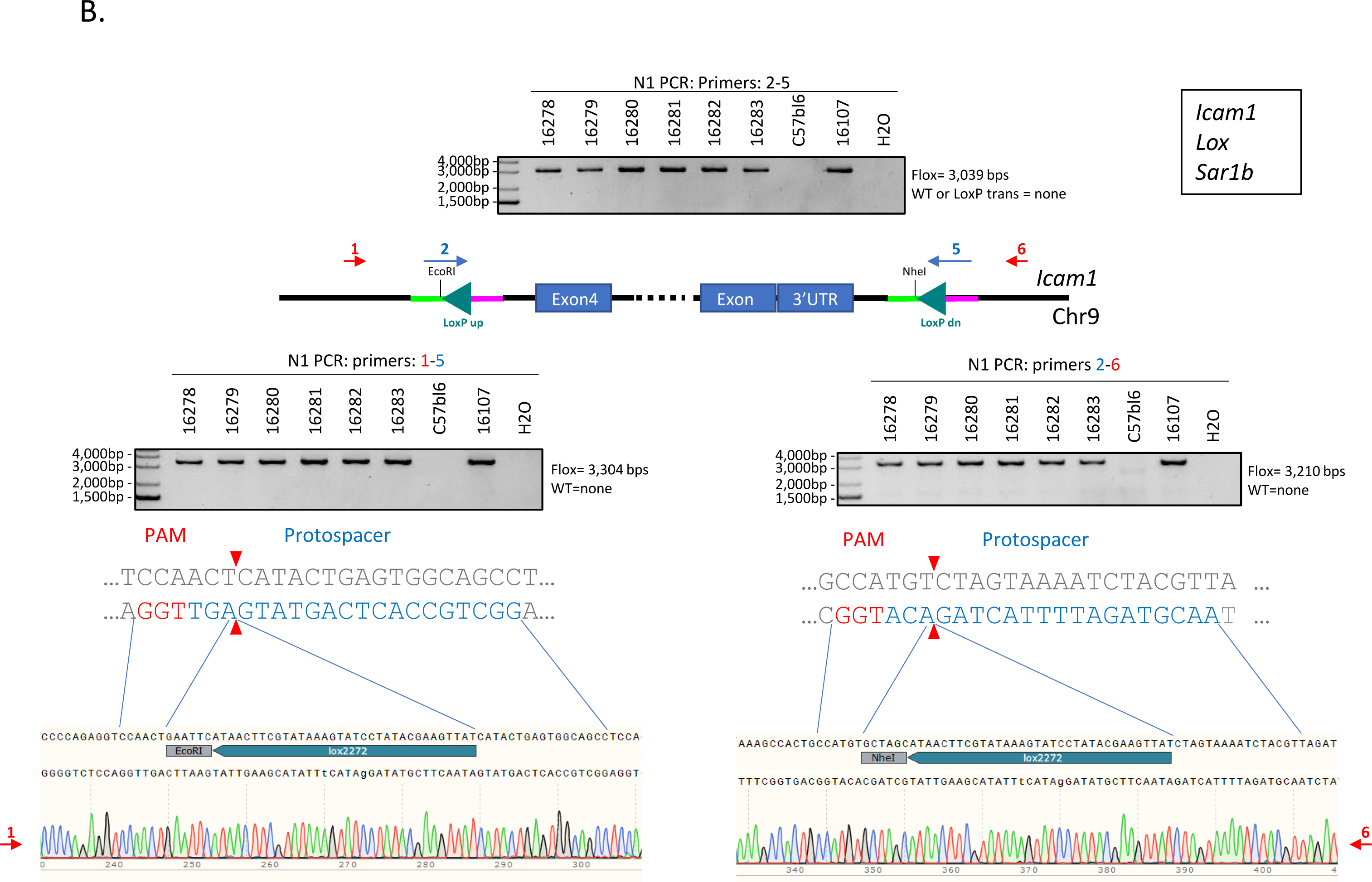
Consecutive sequential electroporation was successfully applied to generate *Icam1* floxed animals. **A**. Schematic representation highlighting the primer positions and genotyping strategy used for *Icam1* floxed F0 animals characterization. PCR results from primers 2-5 combination are depicted in the far-left panel, PCR results from primers 2-6 combination are depicted in the middle panel and PCR results from the primers 1-5 combination are depicted in the far-right panel. **B**. Schematic representation highlighting the primer positions, genotyping strategy and sequencing results obtained from *Icam1* floxed N1 animals. PCR results from primers 2-5 combination are depicted in the upper middle panel, PCR results and sequencing alignment from primers 1-5 combination are depicted in the lower left panel. PCR results and sequencing alignment from primers 2-6 combination are depicted in the lower right panel. Primers used for sequencing are highlighted in red. The same strategy was applied to complete a total of three different projects (*Icam1, Lox, Sar1b*).

Our standard genotyping strategy consisted of using the long *loxP* site overlapping primers 2 and 5 (figure 2A) as an initial screening step to identify any positive animals containing both *loxP* sites in cis (on the same allele). Animals were also investigated by using primers 2-6 and 1-5 combinations in separate PCR reactions (figure 2A). Positive PCR products from these last reactions were then send for sequencing using either primer 1 or 6 depending on the initial primer pairs used (red primers figure 2B). In some instance, primers 3 and 4 were used for further validation. This latest screening strategy was applied to all of the described projects except for the one involving *Pard6g* that required a different approach since it was impossible to obtain a full-length PCR product between the targeted exon due to high sequence complexity (genotyping strategy, see additional file 4). Using this screening method, we were able to recover pups with both *loxP* sites in *cis* for a total of three projects (details see table 5) with an average integration rate of 8% (range from 5 to 13%). Germline transmission was confirmed for two of these projects.

**Table 5:** Details of the projects successfully completed using the consecutive sequential electroporation procedure. The projects completed using the consecutive sequential electroporation procedure are highlighted in the current table. Column 1 highlights the gene name, column 2 highlights the procedure used to complete the project, column 3 highlights the number of embryos electroporated, column 4 highlights the number of embryos implanted, column 5 highlights the number of surgery performed, column 6 highlights the number of gestations obtained, column 7 highlights the number of pups born, column 8 highlights the percentage of live born animals, column 9 highlights the number of properly targeted animals (*loxP* site in cis, confirmed by sequencing), and column 10 highlights the targeting efficiency for each project.

| Gene name | Procedure | Number of embryos electroporated | Number of embryos implanted | Number of surgeries | Number of gestations | Number of pups born | *Percentage of live born animals | Number of pups properly targeted | **Targeting efficiency |
| --- | --- | --- | --- | --- | --- | --- | --- | --- | --- |
| *Icam1* | consecutive sequential electroporation | 309 | 252 | 10 | 7 | 24 | 10% | 2 | 8% |
| *Lox* | consecutive sequential electroporation | 159 | 140 | 5 | 4 | 22 | 16% | 1 | 5% |
| *Sar1b* | consecutive sequential electroporation | 196 | 128 | 5 | 2 | 8 | 6% | 1 | 13% |

**\*Calculated by dividing the number of pups born by the number of implanted embryos**
**\*\*Calculated by dividing the number of properly targeted pups by the total number of pups born**

The remaining five projects were completed using the non-consecutive sequential electroporation strategy. In this case, we focused our investigation on finding positive pups with a single *loxP* site integration on one side and wild-type sequence on the other side (figure 3A). This was achieved by using positive PCR products from the same 2-6 or 1-5 primer pairs described previously and sequencing these PCR products with either primer 1 or 6 depending on the initial primer pairs used (figure 3A). In this case, the sequencing results informed us as to whether or not the insertion site that failed to incorporate the *loxP* site was exempt of indels. If this was the case, an additional primer, overlapping the genomic DNA sequence adjacent to the *loxP* insertion site and a portion of the *loxP* site itself in opposite direction to the one initially designed was used to confirm the integrity of the inserted *loxP* site (primer 7, figure 3A). In this case, PCR products from primers pairs 1-7 and 2-3 were sent for sequencing using primers 1 and 3 respectively. Pups that were exempt of indels in the site that failed to incorporate a *loxP* site on one side and had proper integration of the *loxP* site on the other side were bred for germline transmission. The resulting N1 animals were sequence verified as described above and bred to N2 before being intercrossed to produce embryos that were used to incorporate the missing *loxP* site (Strategy B, figure 1). We reasoned that using this strategy would increase the likelihood to obtain the properly targeted allele as 25% of the embryos would be homozygotes with a single *loxP* site on both allele, 50% would be heterozygotes with a single *loxP* site on one out of two alleles, and 25% would be wild-type. For each project, electroporation on 1-cell stage embryos was performed using the material to incorporate the missing *loxP* site as described above before being implanted in pseudopregnant females (0.5 dpc). The resulting pups were investigated for proper *loxP* targeting as described previously, with priority given to pups showing positive bands using the 2-5 primer pairs (figures 3B, 3C). Using this strategy, we were able to obtain properly targeted pups for the remaining five projects (details see table 6), with a targeting efficiency averaging 11% (ranging from 3 to 25%). Data from the *Loxl1* project were used to compare the targeting efficiency when using ssODNs corresponding to the targeting versus non-targeting strand for insertion of the second *loxP* site (details see table 6). Interestingly, in this case, the ssODN corresponding to the non-targeting strand gave us a greater efficiency, with a value of 6%, when compared to the ssODN corresponding to the targeting strand that only resulted in 3% efficiency. Hence, these results suggest that, for conditional allele model generation using two ssODNs, the choice between using the targeting versus non-targeting strand as a repair template should be determined empirically as repair efficiency using either one of these strands appear to be context dependent. Germline transmission was confirmed in all five projects.

**Table 6:**
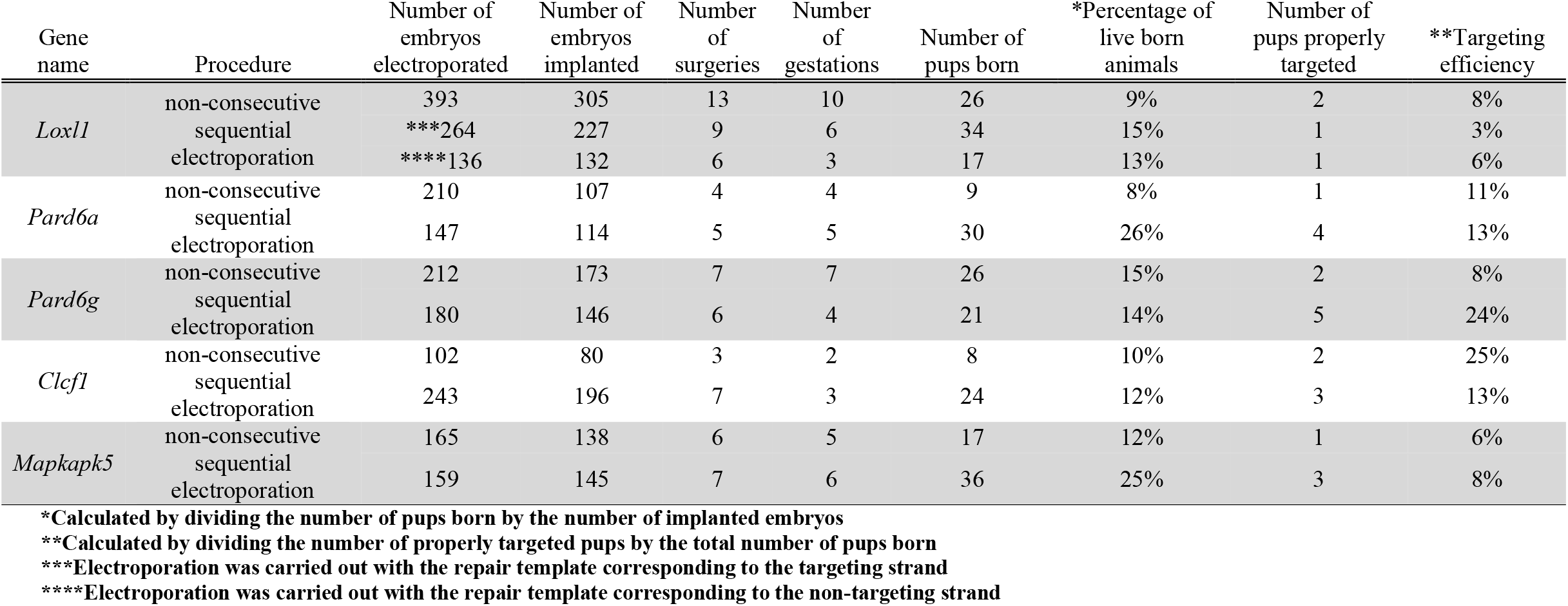
Details of the projects successfully completed using the non-consecutive sequential electroporation procedure. The projects completed using the non-consecutive sequential electroporation procedure are highlighted in the current table. Column 1 highlights the gene name, column 2 highlights the procedure used to complete the project, column 3 highlights the number of embryos electroporated, column 4 highlights the number of embryos implanted, column 5 highlights the number of surgery performed, column 6 highlights the number of gestations obtained, column 7 highlights the number of pups born, column 8 highlights the percentage of live born animals, column 9 highlights the number of properly targeted animals (confirmed by sequencing), and column 10 highlights the targeting efficiency for each project.

**Figure 3:**
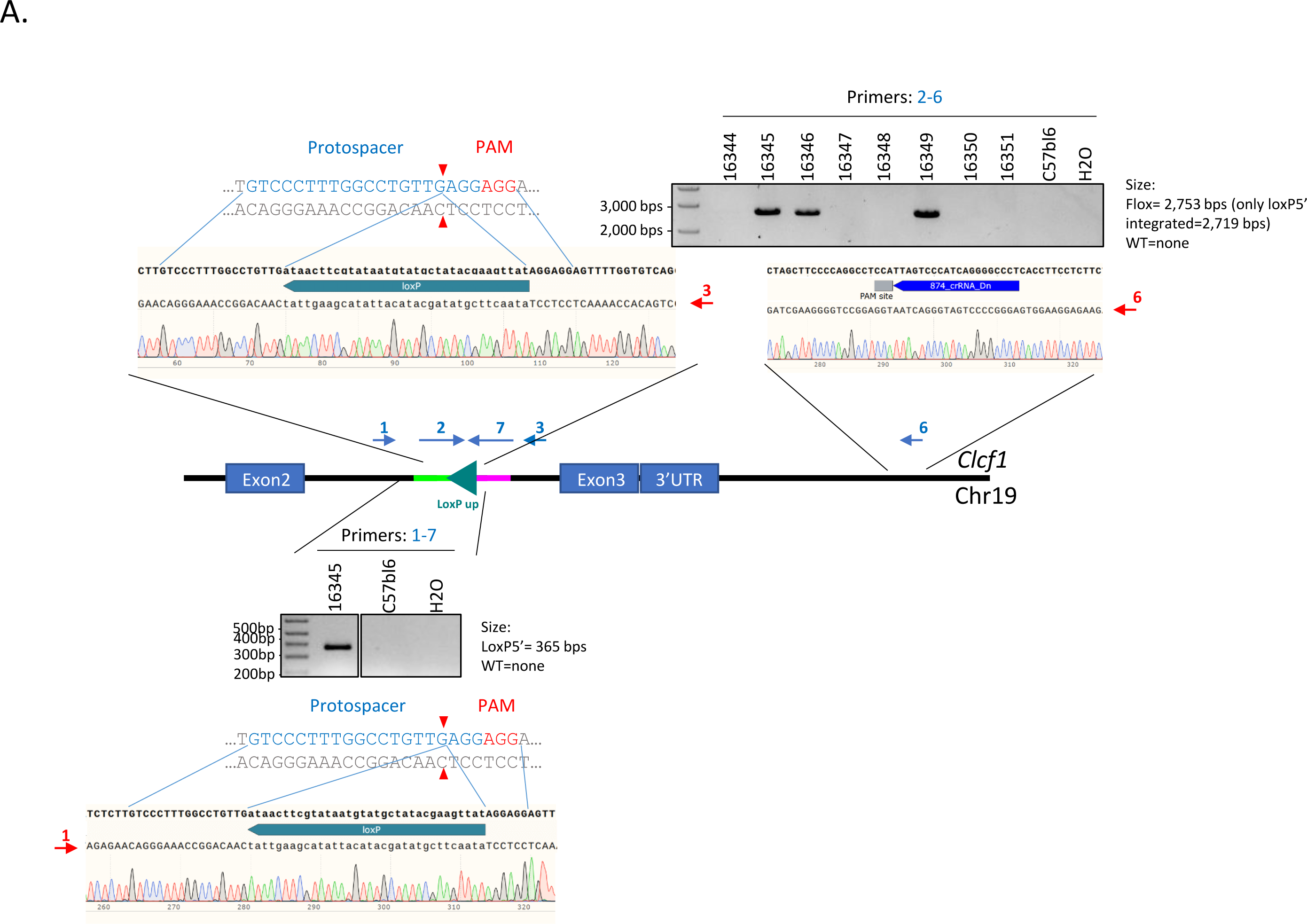
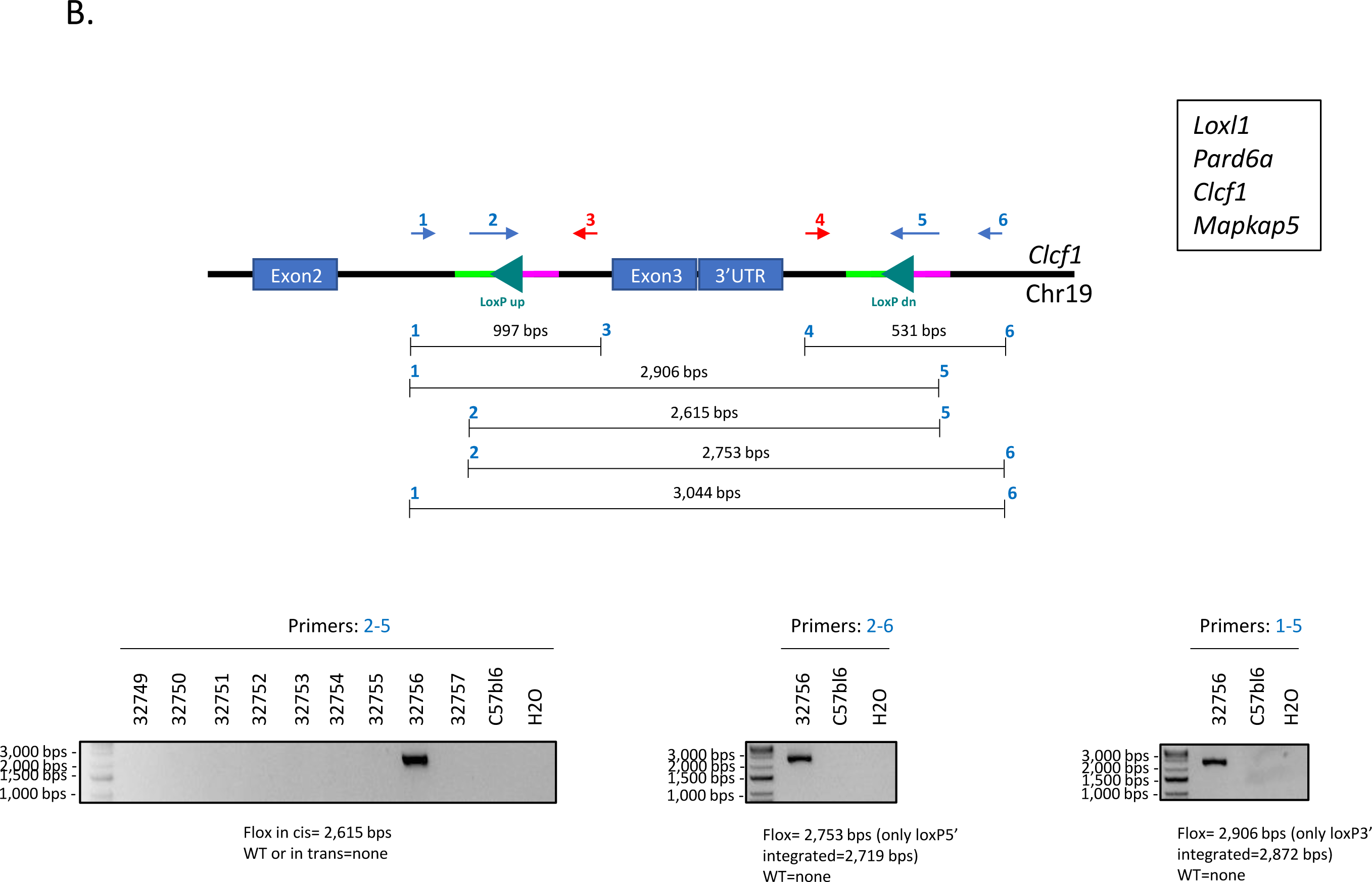
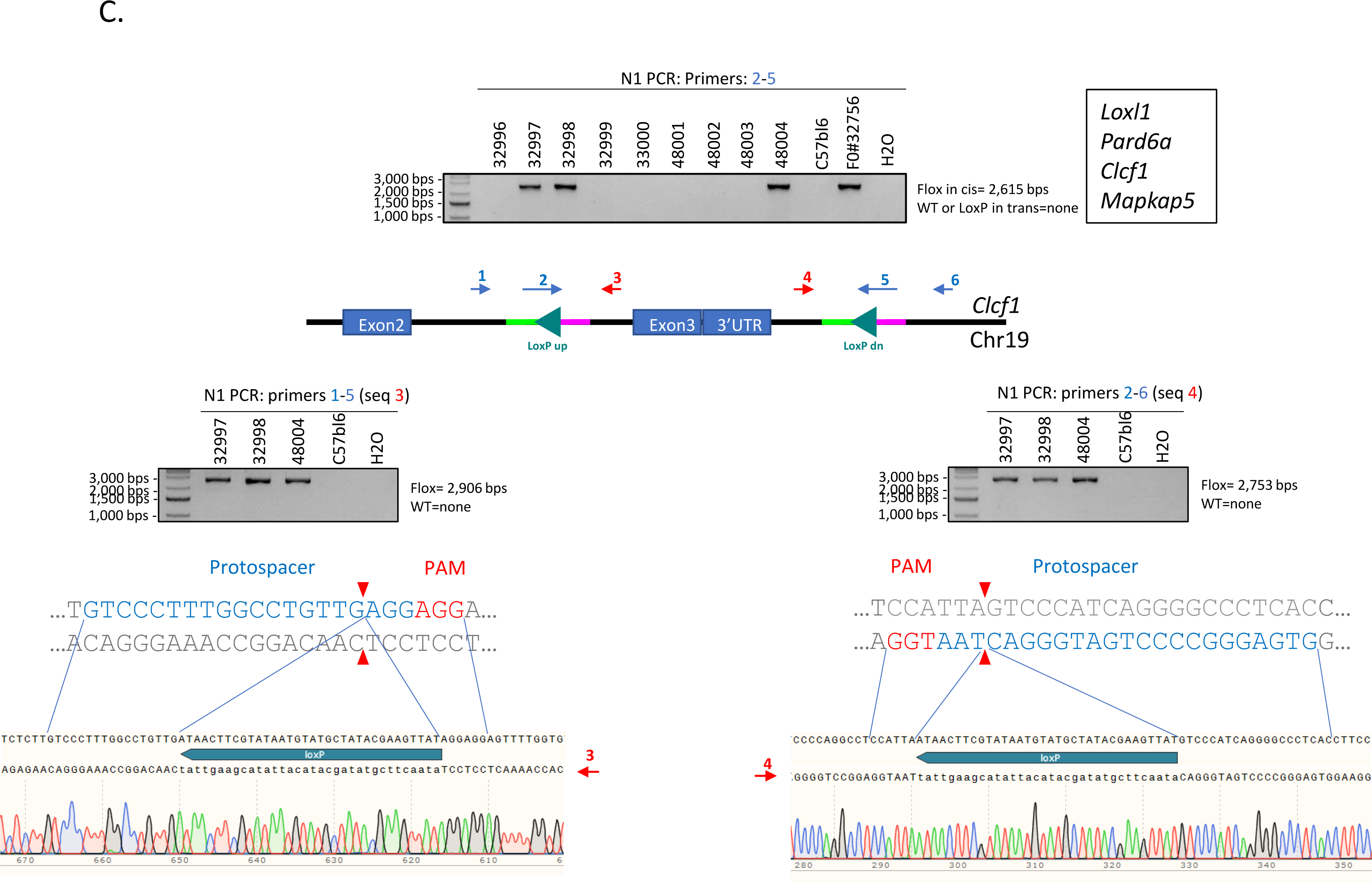
Non-consecutive sequential electroporation was successfully applied to generate *Clcf1* floxed animals. **A**. Schematic representation highlighting the primer positions and genotyping strategy used for *Clcf1* F0 animals characterization. PCR results from primers 2-6 combination are depicted in the upper-right panel. The sequencing results using primers 3 and 6 are highlighted below. PCR results from primers 1-7 combination are depicted in the lower panel. The sequencing results using primer 1 are highlighted below **B**. Schematic representation highlighting the primer positions and genotyping strategy used for *Clcf1* floxed F0 animals characterization. PCR results from primers 2-5 combination are depicted in the far-left panel, PCR results from primers 2-6 combination are depicted in the middle panel and PCR results from the primers 1-5 combination are depicted in the far-right panel. **C**. Schematic representation highlighting the primer positions, genotyping strategy and sequencing results obtained from *Clcf1* floxed N1 animals. PCR results from primers 2-5 combination are depicted in the upper middle panel, PCR results and sequencing alignment from primers 1-5 combination are depicted in the lower left panel. PCR results and sequencing alignment from primers 2-6 combination are depicted in the lower right panel. Primers used for sequencing are highlighted in red. The same strategy was applied to complete a total of four different projects (*Loxl1, Pard6a, Clcf1, Mapkap5*).

## Discussion

The development of novel CRISPR-Cas9 methodologies has improved the efficiency of generating rodent models. Small insertion and exon deletion models are easily generated, however, generating conditional allele models remains a daunting endeavor. Every transgenic core facility functions differently and adapts their methods according to their resources at hand. In our case, all of our services are based on a custom turnkey format; where a researcher come to us with their favorite gene to be targeted and we use our expertise to design the project. We provide the reagents, produce as well as characterize the animal model up to the N1 stage. Hence, to be usable, a method must be flexible, efficient, robust and economical.

Several methods have been described in the literature to generate novel conditional allele models with varying efficiencies. Well established methodologies have relied on double stranded DNA as donor templates requiring extensive homology arms with a targeting efficiency generally reported between 1-10% [2, 16-21]. These repair templates tend to be too long and too complex for simple synthesis, making them less appealing to small platform facilities. The *Easi-*CRISPR method, with its reported efficient targeting (varying between 8.5-100%) and ease of design, was our first method of choice when a large number of conditional allele projects were requested at our facility [7, 8]. However, it became evident that this method could not be applied to all of our projects. Indeed, two of them required targeting constructs outside the 2 kb lssDNA range (Megamer, IDT) and two others targeted regions that were too complex to be synthesized as a lssDNA construct (Megamer, IDT). Furthermore, we faced challenges for two of our ongoing projects with the *Easi*-CRISPR method that resulted in five instances of random construct integration and three other instances of partial construct integration.

These limitations and challenges prompted us to try the sequential electroporation approach reported by Horii *et al*.[10]. We modified the electroporation conditions reported by Troder *et al*., which used a 4 μM Cas9: 4 μM Guide: 10 μM DNA repair template concentration in a single electroporation session [11]. Considering that the Cas9 protein remains active for more than 24 hours after electroporation in embryos, and the fact that 2-cell stage embryos have a similar volume as one cell stage embryos, we rationalized that keeping a final concentration at 4 μM Cas9: 4 μM Guide: 10 μM DNA repair template would be optimal for cleavage efficiency and embryos viability (i.e.: 2 μM: 2 μM; 5 μM on each day). It is noteworthy that the Horii *et al*. publication reported a 20% tetraploidization phenomenon by electrophusion during the second round of electroporation [10]. We did not observe this phenomenon using our electroporation conditions. We hypothesize that this phenomenon is most likely linked to the electroporator used and differences in electroporation conditions. Hence, it is important to keep in mind that the success of the targeting procedure described in our manuscript is highly dependent on the fine tuning of electroporation conditions.

The use of two ssODNs to generate conditional alleles has been the subject of controversy in the transgenic community which is partly due to the difficulties for other groups to reproduce the targeting efficiency obtained in the original manuscript [22, 23]. This appears to be true when using simultaneous injection of all the different components as exemplified by the results from a consortium of 20 transgenic facilities (including ours) reporting a targeting efficiency of less than 1% regardless of the formulation used or delivery method [24]. It is noteworthy that the material used for the *Icam1* project highlighted in this manuscript was also used by our group in the study reported by Gurumurthy *et al*. [24]. Gurumurthy *et al*. also compared the simultaneous injection of two ssODNs to other approaches such as the *Easi-*CRISPR method that was used to complete four different projects, with an average targeting efficiency of ∼13% (ranging from 8 to 18%). These results are comparable to the targeting rate of the consecutive sequential electroporation approach reported in the current manuscript (8%, with a range from 5 to 13%) [24]. Furthermore, the same article reported a method similar to our non-consecutive sequential electroporation procedure; where a first *loxP* site is introduced in embryos by microinjection, and the second *loxP* site is introduced via a second microinjection session with embryos derived from the mouse strain containing the first *loxP* site (referred as second *loxP* site in the next generation) [24, 25]. This method was applied to seven loci that were all successfully flanked with *loxP* sites with a targeting efficiency of 14 ± 6% for the first *loxP* site insertion and 27 ± 32% for the second *loxP* site insertion [24]. Again, these numbers are comparable to our non-consecutive sequential electroporation targeting rate with an average of 11 ± 8% for the first *loxP* site insertion and an average of 11 ± 7% for the second *loxP* site insertion.

In summary, the current manuscript highlights an approach that allows the generation of novel conditional allele models according to two independent possible paths, either via the consecutive or non-consecutive sequential electroporation procedure, with an estimated turnover time varying between 4-10 months depending on the outcome of the initial consecutive sequential electroporation attempt. This method was applied to eight different projects, proving its reliability and flexibility. Furthermore, the fact that this method relies on electroporation rather than microinjection, will certainly be of interest to individual laboratories that do not have access to microinjection set-ups. It is meant to represent an alternative to other already established methods. We also believe that this approach will be of interest to smaller platforms with limited resources for producing large DNA constructs or provide an opportunity to re-visit projects that were not completed using other approaches. One of the main advantages of our method is the relatively inexpensive cost and synthesis turnover time of ssODNs in comparison to lssDNA templates. This provides greater flexibility as more than one guides-repair template combination can be ordered and tested for proper targeting in live animals. This is generally not the case when using a lssDNA template as guide cleavage efficiencies are generally determined in pre-implantation embryos and the repair template is designed based on the two most efficient guides (one in Up and one in Dn position). This approach is based on the assumption that guide cutting efficiency alone dictates the targeting outcome with limited flexibility on the repair template orientation. However, our results suggest that at least for one of our projects, the repair template orientation appears to be critical for targeting efficiency.

New methodologies using the CRISPR-Cas9 tool are being published constantly. These new methodologies allow us to circumvent some of the limitations initially observed for some of the projects highlighted in this manuscript. One of these limitations is the size limit for synthesis of commercially available lssDNA that was limited to 2 kb (Megamer, IDT). It has been recently reported that lssDNA up to 3.5 kb spanning repetitive sequences have been developed using the CRISPR-Clipped *Long* ssDNA via *I*ncising *P*lasmid (CRISPR-CLIP) approach [26]. Although interesting, this method still depends on the establishment of the CRISPR-CLIP approach in an individual laboratory which represents as substantial investment in comparison to the ease and flexibility of ordering short ssODN from commercially available vendors. Furthermore, two commercial vendors, Genscript and Genwiz are now advertising lssDNA production of up to 4 and 10 kb respectively. These new possibilities allow to overcome the previously mentioned size limit, but also represents a substantial monetary investment in comparison to the cost of short ssODN. Embryo electroporation is becoming routinely used in transgenic core facilities as it allows for the manipulation of large numbers of embryos with relatively high editing efficiency. It was recently demonstrated that generating conditional allele models by electroporating two ssODNs *in utero* was possible. This method, streamlines the current procedure by bypassing the *ex vivo* embryo handling steps [27]. This report, although exciting, has reported the generation of properly targeted animals in hybrid mouse strains only, which could be considered a limitation when taking into consideration the number of backcrosses that would be required to bring these targeted alleles onto a pure background. Embryos of inbred strains were demonstrated to be properly targeted using this approach, however, it remains to be proven that this method could results in properly targeted live animals before considering its usage in transgenic facilities. Finally, we note that a recent paper has shown that the addition of RAD51 purified protein along with the CRISPR-Cas9 reagents can significantly increase homozygous knockin in mouse embryos [28]. Hence, it would be interesting to investigate whether or not this homozygous knockin improvement has any influence on cis *loxP* targeting rate in our consecutive sequential electroporation procedure.

## Conclusions

In this work, we have refined the sequential electroporation procedure described by Horii *et al*. resulting in the production of conditional allele models for eight different genes via two different possible paths. We believe that our strategies, demonstrated to be reproducible for eight different loci, should be considered as an alternative to other well-established methodologies in the literature for conditional allele mouse model generation.

## Methods

### Animals

C57BL/6N embryos were produced from males and females purchased from Charles River laboratories, whereas C57BL/6J embryos were produced from males and females purchased from Jackson Laboratory. Hsd:ICR (CD-1) female mice from Envigo+ were used for embryo transfers. All mice were maintained in the pathogen-free Centre de Recherche du Centre Hospitalier de l’Université de Montréal (CRCHUM) animal facility on a 6:30 AM to 6:30 PM light cycle, 21-26°C with 40-60% relative humidity, and had food and water *ad libitum*.

### Guide selection process

Literature reviews and gene structure analyses were performed for each project. When appropriate, the same exons used in previously published classical models were selected for *loxP* sites integration. The ATG containing exons and exons inducing frameshifts were preferentially selected. When possible, exons including 5’ and 3’ regulatory regions were avoided. Once the selection process was completed, *in silico* guide cutting surveys were conducted on genomic DNA located at least 100 base pairs away from the selected exons. Surveys were conducted with the three-following software: CRISPOR (http://crispor.tefor.net/crispor.py), CHOPCHOP (https://chopchop.cbu.uib.no/) and Breaking-Cas (https://bioinfogp.cnb.csic.es/tools/breakingcas/) [13-15]. Guide with good predicted cleavage activity in all three software and falling within regions containing low sequence complexity were retained for the procedure.

### CRISPR-Cas9 Reagents

The Cas9 protein (Integrated DNA technologies (IDT, catalog number 1081058), custom crRNA (IDT, Alt-R™ crRNA) and generic tracrRNA (IDT, catalog number 1072533) were prepared as previously described [11, 29]. Briefly, 50 uM crRNA-tracrRNA annealed complex (pgRNA) were formed by mixing an equimolar ratio of each component that were incubated 5 minutes at 95°C and allowed to cool down to room temperature for 10 minutes. A complete list of guides used for each project is detailed in table 3.

### Preparation of CRISPR-Cas9 electroporation mixes

The Cas9 RNP complex was assembled as previously described [11, 29]. Briefly, 40 μmoles of Cas9 protein (IDT, 1081058) was incubated with 40 μmoles of assembled pgRNA and incubated for 10 minutes at room temperature. The RNP complex was combined in 20 μl Opti-MEM at a final concentration of 2 μM (ThermoFisher Scientific catalog number 31985070) along with 5 μM of repair template (custom Ultramer ssDNA, IDT). Repair template details are highlighted in table 2.

### Zygote preparation

Prepubescent 3 weeks old C57BL/6N or C57BL/6J females were superovulated 67 hours prior zygote collection by 5 IU intraperitoneal injection of pregnant mare serum gonadotrophin (Genway Biotech Inc, GWB-2AE30A) followed 47-48 hours later with 5 IU of human chorionic gonadotrophin (Sigma-Aldrich, CG10-1VL) before being bred. Fertilized 1-cell stage embryos were collected and kept in embryomax KSOM advance media (Millipore Sigma cat number MR-101-D) at 37°C under 5% CO_2_ until electroporation (around 12:30 to 2:30 pm).

### Consecutive sequential electroporation procedure

Briefly, 1-cell embryos were washed in batch of 50 through 5 drops of M2 media before being washed in a single drop of Opti-MEM. The embryos were transferred to the 20 ul first Cas9-RNP-ssODN mix. The solution was transferred to a pre-warmed 1 mm cuvette (BioRad). Electroporation was carried out using a Gene Pulser XCell electroporator with the following conditions: 30V, 3 ms pulse duration, 2 pulse 100 ms interval. Electroporated embryos were flush recovered from the cuvette and washed in three drops of embryomax KSOM advance media before being incubated overnight at 37°C under 5% CO_2_. Embryos that developed to the 2-cell stage were subsequently electroporated with the second Cas9-RNP-ssODN mix as described above. These 2-cell electroporated embryos were recovered and washed in embryomax KSOM advance media before being incubated at least one hour at 37°C under 5% CO_2_ prior implantation in pseudopregnant females (0.5 dpc).

### Non-consecutive sequential electroporation procedure

For non-consecutive sequential electroporation procedure embryos were collected from intercross between N2 males and females with one properly targeted *loxP* site. 1-cell embryos collection and electroporation were performed as described in the consecutive sequential electroporation section. RNP complexes and repair template concentrations were adjusted to 4 μM and 10 μM respectively. 1-cell stage electroporated embryos were incubated at least one hour at 37°C under 5% CO_2_ prior implantation in pseudopregnant females (0.5 dpc).

### Genotyping

Ear biopsies from 21 days old mice were digested in MyTaq Extract PCR kit according to the manufacturer (Bioline, Cat number BIO-21126). PCR amplification was performed using the high-fidelity enzyme Q5 from New England Biolab (NEB, Cat number M0530L) or the Platinum SuperFi DNA polymerase from ThermoFisher (Cat number 123551010) for hard to amplify locus (*Lox*). Amplification was performed with locus specific primers detailed in table 4 [8]. The PCR products obtained were sent for sequencing at the Centre d’expertise et de service de génome Québec (CHU-Ste-Justine, Montréal, Canada). Sequence alignments were analyzed using the Snapgene software (https://www.snapgene.com/).

### *In vitro* Cre recombination assay

Ear biopsies from 21 days old mice of the appropriate genotype were digested using the DNeasy Blood and Tissue Kits from Qiagen (Cat number 69504). The purified DNA was quantified using a Nanodrop spectrometer and the DNA concentration for each sample was adjusted to 35 ng/μl. A volume of 5 μl of genomic DNA (175 ng) in a total reaction volume of 25 μl, including 1-2 units of NEB Cre recombinase (Cat number M0298S) and its supplied buffer was incubated overnight at 37°C. A volume of 3.5 μl of the Cre reaction was then used for PCR amplification using the high-fidelity enzyme Q5 (NEB, M0530L) as described above.

## List of abbreviation

NHEJ: Non-homologous end joining
HDR: Homology dependent repair
Indels: insertions and deletions
lssDNA: long single stranded DNA
CLICK: CRISPR with lssDNA inducing conditional knockout allele
ssODN: short single strand oligonucleotide
CRISPR-CLIP: CRISPR-Clipped *Long* ssDNA via *I*ncising *P*lasmid
CRCHUM: Centre de Recherche du Centre Hospitalier de l’Université de Montréal

## Declaration

### Ethics approval and consent to participate

All animal care and procedures performed in this study were approved by the CRCHUM animal care committee in accordance with the guidelines from the Canadian Council on Animal Care in science (CCAC); protocol number N17026JFSrs.

### Consent for publication

Not applicable

### Availability of data and materials

The datasets used and/or analysed during the current study are available from the corresponding author on reasonable request.

### Authors’ contributions

GB and JFS designed the projects. HJ, CL and EL contributed to reagents. GB, MO, AB, and JFS contributed to the electroporation sessions, mouse models generation and characterization process. GB contributed to the figures and data interpretations. JFS performed all data analyses and wrote the manuscript.

## Acknowledgements

We thank the principal investigators that agreed on sharing the data related to their specific projects. We also thank Jon Neumann from UC Irvine for sharing the *in vitro* Cre recombination assay as well as Dre. Christine Vande Velde, and Dr. Greg Fitzharris for aid in manuscript preparation.

