## Additional file 1 for "Introduction of *loxP* sites by electroporation in the mouse genome; a simple approach for conditional allele generation in complex targeting loci"

**Additional files**

| Gene name | Procedure | Number of embryos microinjected | Number of embryos that survived | Number of embryos implanted | Concentration (ng/μl) | Number of surgeries | Number of gestations | Number of pups born | Number of properly targeted pups | Random integration | Partial integration |
| --- | --- | --- | --- | --- | --- | --- | --- | --- | --- | --- | --- |
| *Sar1b* | *Easi*-CRISPR | 413 | 334 | 297 | 10:10:10-15:15:10 | 11 | 7 | 18 | 0 | 3 | 1 |
| *Loxl1* | *Easi*-CRISPR | 593 | 487 | 448 | 5:5:5-20:20:20 | 17 | 12 | 25 | 0 | 2 | 2 |

**Additional file 1: Details of the projects that were not completed using the *Easi*-CRISPR procedure.**

The projects that were not completed using the *Easi*-CRISPR procedure are highlighted in the current table. Column 1 highlights the gene name, column 2 highlights the procedure used, column 3 highlights the number of embryos microinjected, column 4 highlights the number of embryos that survived the microinjection, column 5 highlights the number of embryos implanted, column 6 highlights the reagent concentrations used (Cas9:Guide RNA:repair template), column 7 highlights the number of surgeries performed, column 8 highlights the number of gestations obtained, column 9 highlights the number of pups born, column 10 highlights the number of properly targeted animals, column 11 highlights the number of pups containing random construct integrations and column 12 highlights the number of pups containing partial construct integrations at the targeted site for each project.

**Additional file 2: The *Easi-*CRISPR procedure resulted in random and partial *Sar1b* floxed construct integration**

**A.** Schematic representation and corresponding DNA sequence highlighting the repair template and primer positions for *Sar1b* floxed animals generation using *Easi*-CRISPR. Corresponding flanking sequence are in black, 5’homology arm sequence is in green, 3’homology arm sequence is in pink, intronic region sequences are in yellow, coding region sequences are in blue, *loxP* corresponding sequences are in dark green. Blue arrows highlight genotyping primer sequences. **B.** Schematic representation highlighting primer positions and genotyping strategy used for *Sar1b* floxed F0 animals characterization using *Easi*-CRISPR. PCR results from primers 1-6 combination are depicted in the upper-left panel, PCR results from primers 2-5 combination are depicted in the lower-left panel, PCR results from primers 1-5 combination are depicted in the middle panel, and PCR results from primers 2-6 combination are depicted in the far-right panel. *Highlights animals containing random construct integration.

**Additional file 3: The *Easi-*CRISPR procedure resulted in random and partial *Loxl1* floxed construct integration**

**A.** Schematic representation and corresponding DNA sequence highlighting the repair template and primer positions for *Loxl1* floxed animals generation using *Easi*-CRISPR. Corresponding flanking sequence are in black, 5’homology arm sequence is in green, 3’homology arm sequence is in pink, intronic region sequences are in yellow, coding region sequences are in blue, *loxP* corresponding sequences are in dark green. Blue arrows highlight genotyping primer sequences. **B.** Schematic representation highlighting the primer positions and genotyping strategy used for *Loxl1* floxed F0 animals characterization using *Easi*-CRISPR. PCR results from primers 1-6 combination are depicted in the upper-left panel, PCR results from primers 2-5 combination are depicted in the lower-left panel, PCR results from primers 1-5 combination are depicted in the middle panel, and PCR results from primers 2-6 combination are depicted in the far-right panel. *Highlights animals containing random construct integration. **Highlights animals with partial construct integration at the targeted site.

**Additional file 4: Non-consecutive sequential electroporation was successfully applied to generate *Pard6g* floxed animals**

**A.** Schematic representation highlighting the primer positions and genotyping strategy used for *Pard6g* floxed F0 animals characterization. PCR and sequencing results from primers 1-7 combination are depicted in the far-left panel, PCR and sequencing results from primers 2-3 combination are depicted in the middle-left panel, PCR and sequencing results from primers 4-5 combination are depicted in the middle-right panel, PCR and sequencing results from primers 8-6 combination are depicted in the far-right panel. Primers used for sequencing are highlighted in red. **B.** Schematic representation illustrating primer positions and PCR products resulting from *in vitro* Cre recombination of the *Pard6g* locus. PCR results amplification using primers 1-6 combination from purified gDNA untreated or treated with Cre-recombinase are depicted in the lower panels with their expected molecular weight. The difference in Cre+ molecular weight is due to the presence of two different *loxP* sites in the Up position. The Up1 *loxP* site is positioned 30 base pairs farther than the Up3 *loxP* site. Cre-, untreated gDNA; Cre+, treated gDNA.
