## Additional file 2 for "Introduction of *loxP* sites by electroporation in the mouse genome; a simple approach for conditional allele generation in complex targeting loci"

### Slide 1
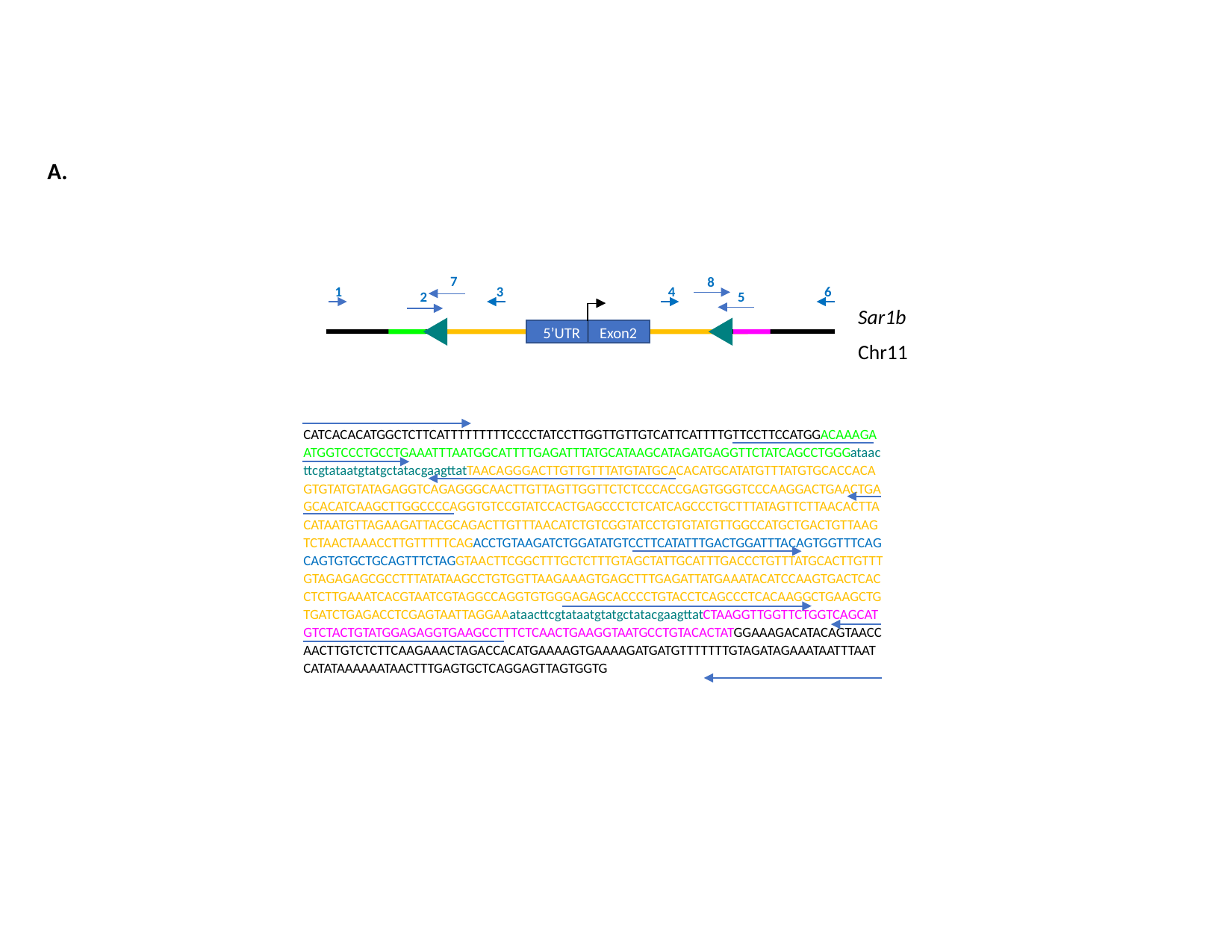

A.
7
8
1
3
4
6
2
5
5’UTR
Exon2
Sar1b
Chr11
CATCACACATGGCTCTTCATTTTTTTTTCCCCTATCCTTGGTTGTTGTCATTCATTTTGTTCCTTCCATGGACAAAGAATGGTCCCTGCCTGAAATTTAATGGCATTTTGAGATTTATGCATAAGCATAGATGAGGTTCTATCAGCCTGGGataacttcgtataatgtatgctatacgaagttatTAACAGGGACTTGTTGTTTATGTATGCACACATGCATATGTTTATGTGCACCACAGTGTATGTATAGAGGTCAGAGGGCAACTTGTTAGTTGGTTCTCTCCCACCGAGTGGGTCCCAAGGACTGAACTGAGCACATCAAGCTTGGCCCCAGGTGTCCGTATCCACTGAGCCCTCTCATCAGCCCTGCTTTATAGTTCTTAACACTTACATAATGTTAGAAGATTACGCAGACTTGTTTAACATCTGTCGGTATCCTGTGTATGTTGGCCATGCTGACTGTTAAGTCTAACTAAACCTTGTTTTTCAGACCTGTAAGATCTGGATATGTCCTTCATATTTGACTGGATTTACAGTGGTTTCAGCAGTGTGCTGCAGTTTCTAGGTAACTTCGGCTTTGCTCTTTGTAGCTATTGCATTTGACCCTGTTTATGCACTTGTTTGTAGAGAGCGCCTTTATATAAGCCTGTGGTTAAGAAAGTGAGCTTTGAGATTATGAAATACATCCAAGTGACTCACCTCTTGAAATCACGTAATCGTAGGCCAGGTGTGGGAGAGCACCCCTGTACCTCAGCCCTCACAAGGCTGAAGCTGTGATCTGAGACCTCGAGTAATTAGGAAataacttcgtataatgtatgctatacgaagttatCTAAGGTTGGTTCTGGTCAGCATGTCTACTGTATGGAGAGGTGAAGCCTTTCTCAACTGAAGGTAATGCCTGTACACTATGGAAAGACATACAGTAACCAACTTGTCTCTTCAAGAAACTAGACCACATGAAAAGTGAAAAGATGATGTTTTTTTGTAGATAGAAATAATTTAATCATATAAAAAATAACTTTGAGTGCTCAGGAGTTAGTGGTG

### Slide 2
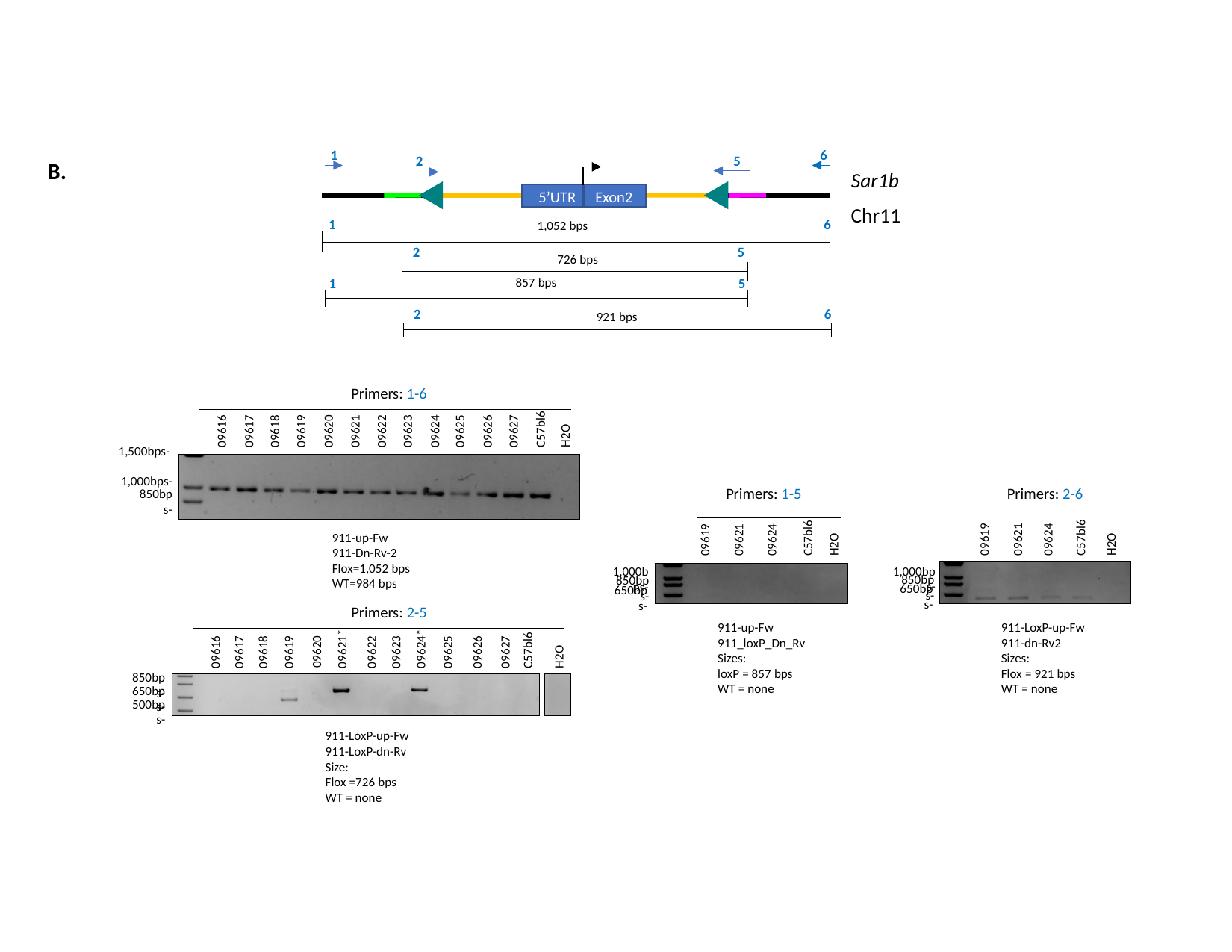

1
6
2
5
Sar1b
5’UTR
Exon2
Chr11
1
6
1,052 bps
2
5
726 bps
1
5
857 bps
2
6
921 bps
Primers: 1-6
09621
09622
09623
09624
09625
09626
09627
C57bl6
09616
09617
09618
09619
09620
H2O
1,500bps-
1,000bps-
850bps-
Primers: 1-5
Primers: 2-6
09619
09621
09624
C57bl6
H2O
911-up-Fw
911-Dn-Rv-2 Flox=1,052 bps
WT=984 bps
09619
09621
09624
C57bl6
H2O
1,000bps-
1,000bps-
850bps-
850bps-
650bps-
650bps-
Primers: 2-5
911-up-Fw
911_loxP_Dn_Rv
Sizes:
loxP = 857 bps
WT = none
911-LoxP-up-Fw
911-dn-Rv2
Sizes:
Flox = 921 bps
WT = none
09616
09617
09618
09619
09621*
09622
09623
09624*
09625
09626
09627
C57bl6
H2O
09620
850bps-
650bps-
500bps-
911-LoxP-up-Fw
911-LoxP-dn-Rv
Size:
Flox =726 bps
WT = none
B.
