## Additional file 3 for "Introduction of *loxP* sites by electroporation in the mouse genome; a simple approach for conditional allele generation in complex targeting loci"

### Slide 1
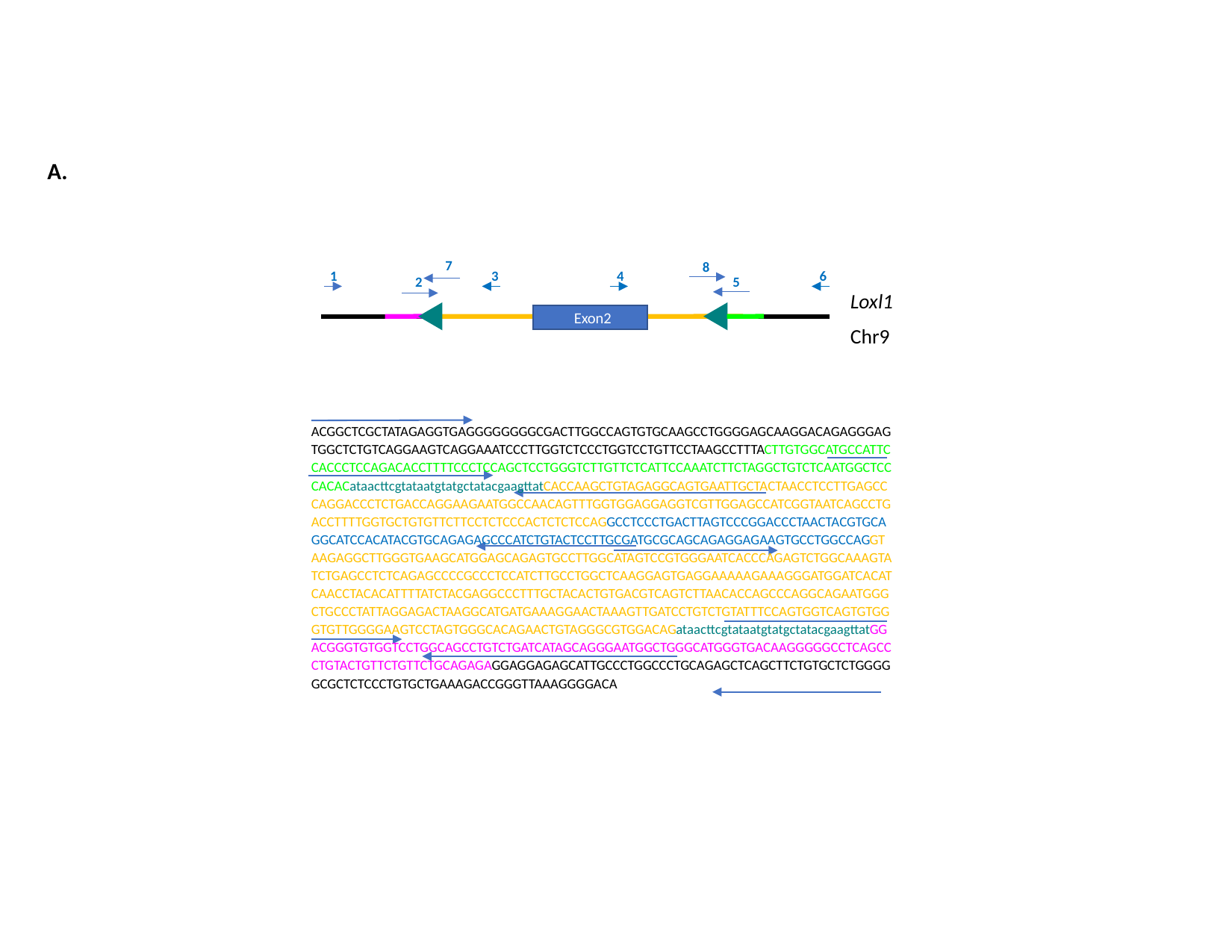

A.
7
8
1
3
4
6
2
5
Exon2
Loxl1
Chr9
ACGGCTCGCTATAGAGGTGAGGGGGGGGCGACTTGGCCAGTGTGCAAGCCTGGGGAGCAAGGACAGAGGGAGTGGCTCTGTCAGGAAGTCAGGAAATCCCTTGGTCTCCCTGGTCCTGTTCCTAAGCCTTTACTTGTGGCATGCCATTCCACCCTCCAGACACCTTTTCCCTCCAGCTCCTGGGTCTTGTTCTCATTCCAAATCTTCTAGGCTGTCTCAATGGCTCCCACACataacttcgtataatgtatgctatacgaagttatCACCAAGCTGTAGAGGCAGTGAATTGCTACTAACCTCCTTGAGCCCAGGACCCTCTGACCAGGAAGAATGGCCAACAGTTTGGTGGAGGAGGTCGTTGGAGCCATCGGTAATCAGCCTGACCTTTTGGTGCTGTGTTCTTCCTCTCCCACTCTCTCCAGgcctccctgacttagtcccggaccctaactacgtgcaggcatccacatacgtgcagagagcccatctgtactccttgcgatgcgcagcagaggagaagtgcctggccagGTAAGAGGCTTGGGTGAAGCATGGAGCAGAGTGCCTTGGCATAGTCCGTGGGAATCACCCAGAGTCTGGCAAAGTATCTGAGCCTCTCAGAGCCCCGCCCTCCATCTTGCCTGGCTCAAGGAGTGAGGAAAAAGAAAGGGATGGATCACATCAACCTACACATTTTATCTACGAGGCCCTTTGCTACACTGTGACGTCAGTCTTAACACCAGCCCAGGCAGAATGGGCTGCCCTATTAGGAGACTAAGGCATGATGAAAGGAACTAAAGTTGATCCTGTCTGTATTTCCAGTGGTCAGTGTGGGTGTTGGGGAAGTCCTAGTGGGCACAGAACTGTAGGGCGTGGACAGataacttcgtataatgtatgctatacgaagttatGGACGGGTGTGGTCCTGGCAGCCTGTCTGATCATAGCAGGGAATGGCTGGGCATGGGTGACAAGGGGGCCTCAGCCCTGTACTGTTCTGTTCTGCAGAGAGGAGGAGAGCATTGCCCTGGCCCTGCAGAGCTCAGCTTCTGTGCTCTGGGGGCGCTCTCCCTGTGCTGAAAGACCGGGTTAAAGGGGACA

### Slide 2
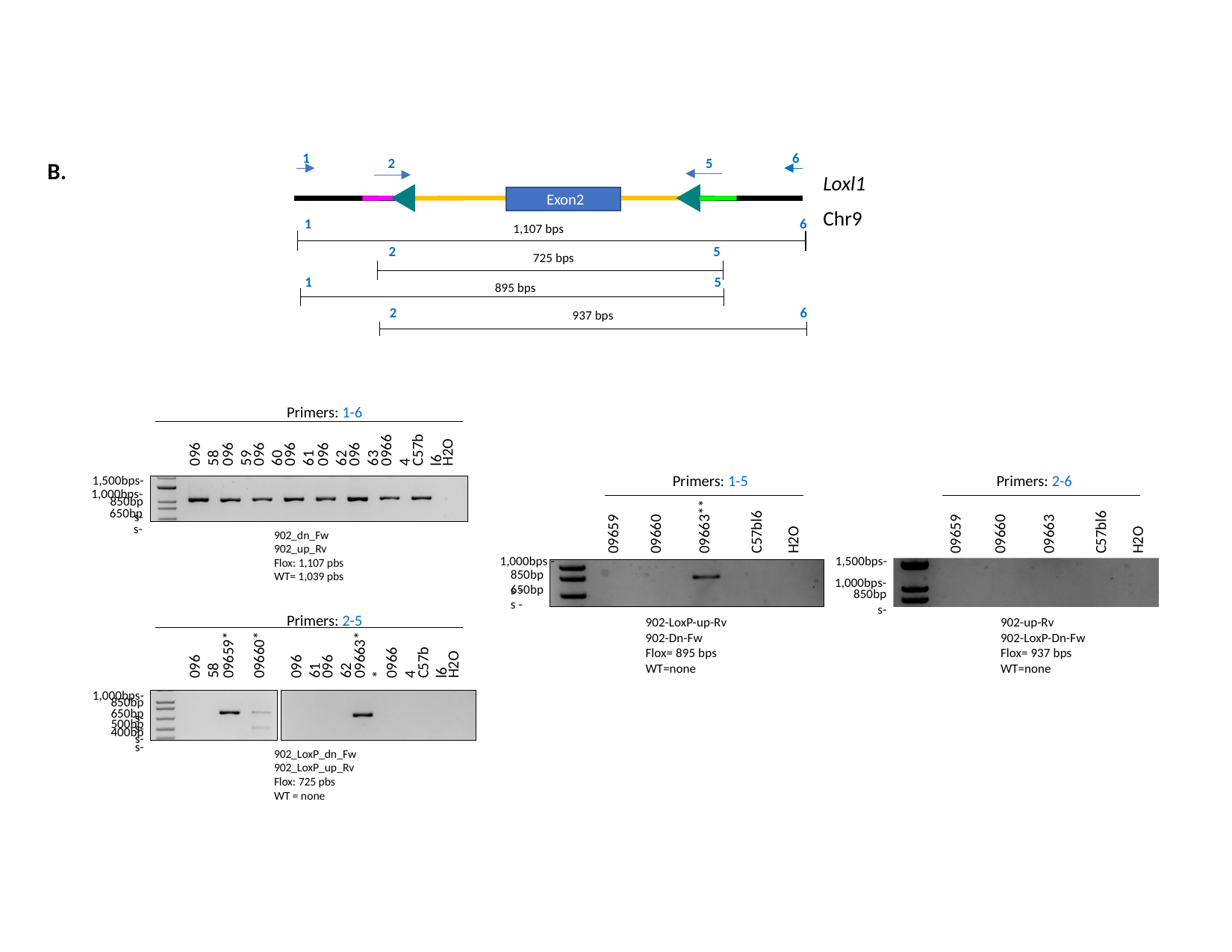

1
6
2
5
Loxl1
Exon2
Chr9
1
6
1,107 bps
2
5
725 bps
1
5
895 bps
2
6
937 bps
Primers: 1-6
H2O
09664
C57bl6
09658
09659
09660
09661
09662
09663
1,500bps-
1,000bps-
Primers: 1-5
Primers: 2-6
850bps-
650bps-
09663**
09660
C57bl6
09660
C57bl6
09659
H2O
09659
09663
H2O
902_dn_Fw 902_up_Rv
Flox: 1,107 pbs
WT= 1,039 pbs
1,000bps -
1,500bps-
1,000bps-
850bps -
650bps -
850bps-
Primers: 2-5
902-LoxP-up-Rv
902-Dn-Fw
Flox= 895 bps
WT=none
902-up-Rv
902-LoxP-Dn-Fw
Flox= 937 bps
WT=none
09659*
09660*
09663**
H2O
09664
C57bl6
09658
09661
09662
1,000bps-
850bps-
650bps-
500bps-
400bps-
902_LoxP_dn_Fw
902_LoxP_up_Rv
Flox: 725 pbs
WT = none
B.
