## Additional file 4 for "Introduction of *loxP* sites by electroporation in the mouse genome; a simple approach for conditional allele generation in complex targeting loci"

### Slide 1
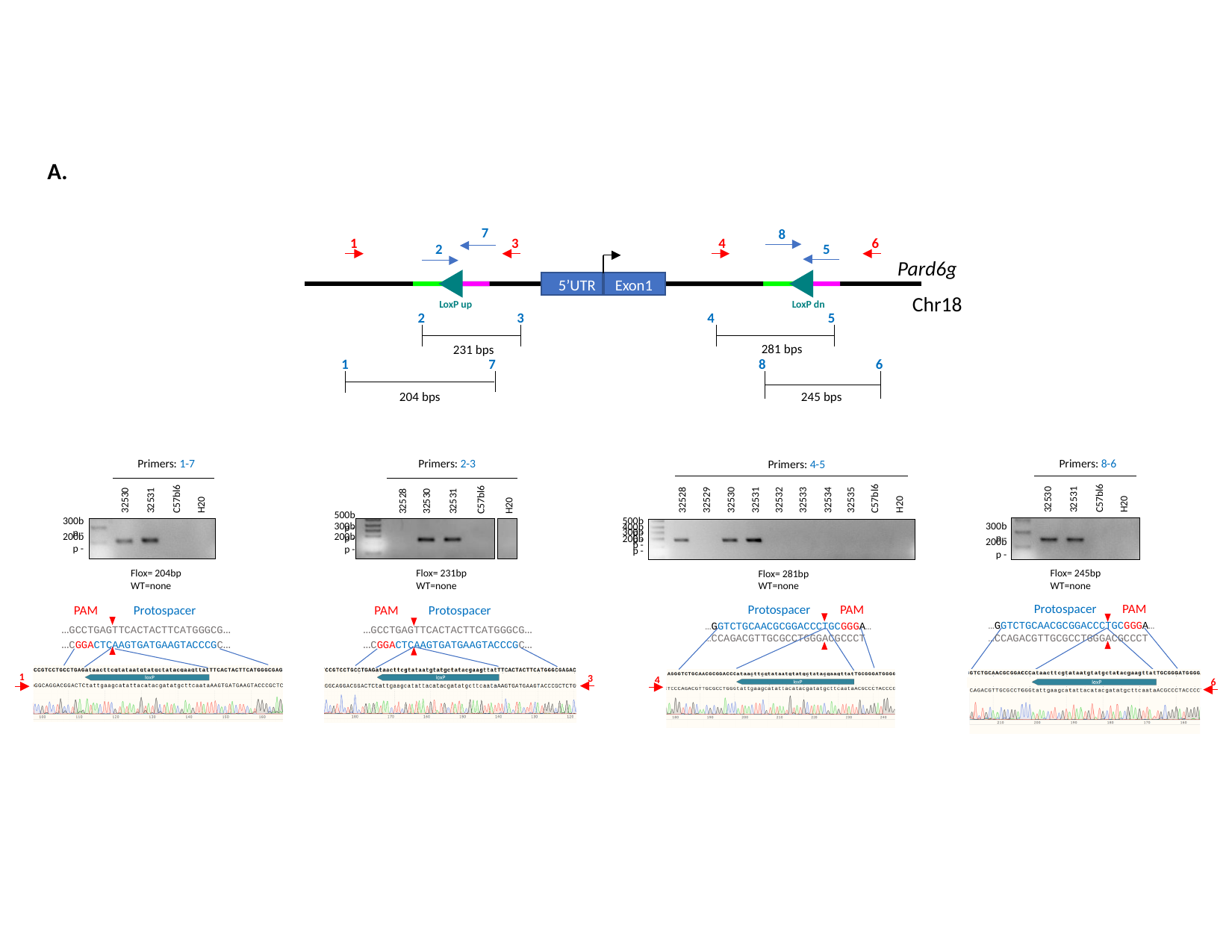

A.
7
8
1
3
4
6
2
5
5’UTR
Exon1
LoxP up
LoxP dn
2
3
4
5
281 bps
231 bps
1
7
8
6
245 bps
204 bps
Pard6g
Chr18
Primers: 1-7
Primers: 2-3
Primers: 8-6
Primers: 4-5
32530
32531
C57bl6
H20
32528
32529
32530
32531
32532
32533
32534
32535
C57bl6
H20
32530
32531
C57bl6
H20
32528
32530
32531
C57bl6
H20
500bp -
300bp -
500bp -
300bp -
300bp -
400bp -
300bp -
200bp -
200bp -
200bp -
200bp -
Flox= 204bp
WT=none
Flox= 231bp
WT=none
Flox= 245bp
WT=none
Flox= 281bp
WT=none
Protospacer
PAM
Protospacer
PAM
PAM
Protospacer
PAM
Protospacer
…GGTCTGCAACGCGGACCCTGCGGGA…
…CCAGACGTTGCGCCTGGGACGCCCT
…GGTCTGCAACGCGGACCCTGCGGGA…
…CCAGACGTTGCGCCTGGGACGCCCT
…GCCTGAGTTCACTACTTCATGGGCG…
…CGGACTCAAGTGATGAAGTACCCGC…
…GCCTGAGTTCACTACTTCATGGGCG…
…CGGACTCAAGTGATGAAGTACCCGC…
1
3
4
6

### Slide 2
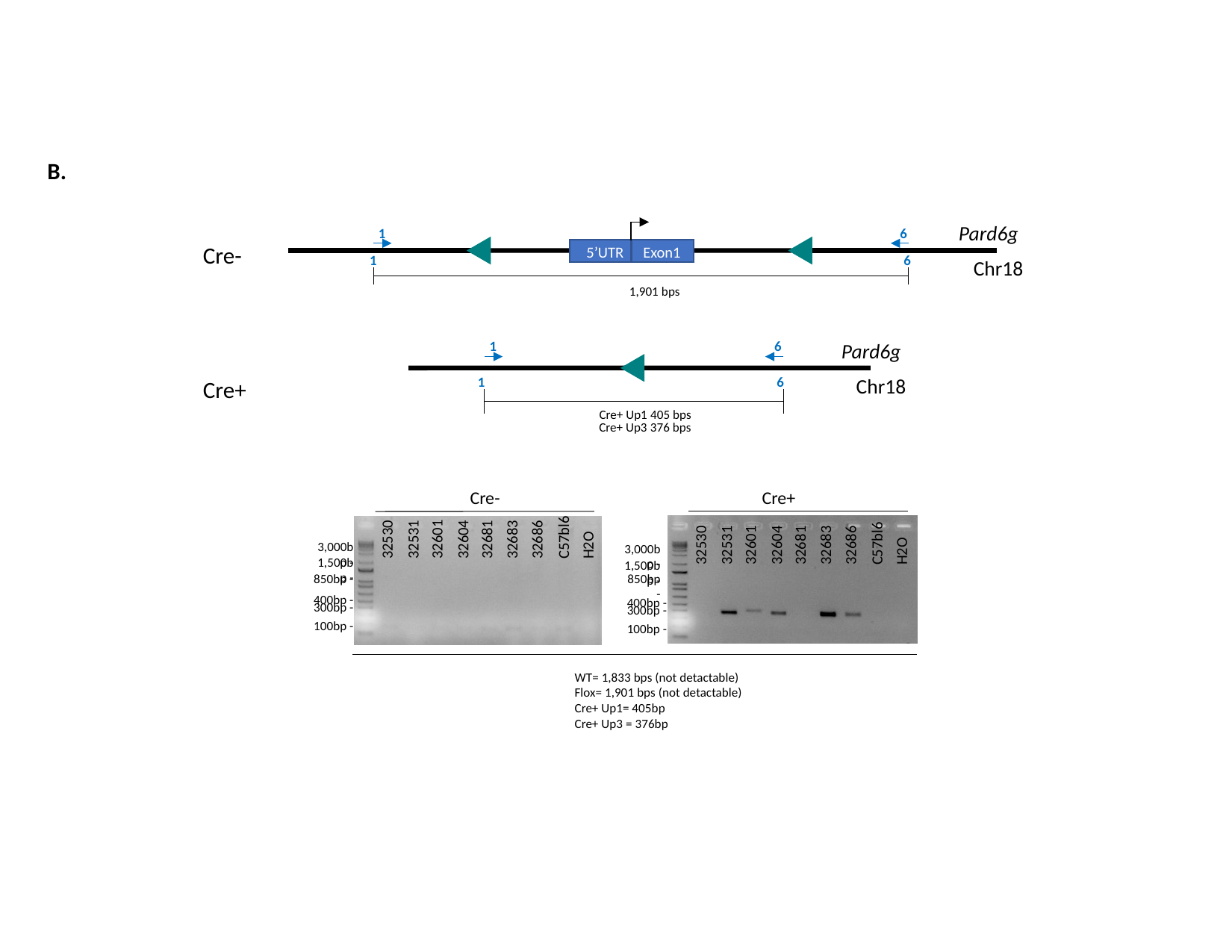

B.
Pard6g
1
6
Cre-
5’UTR
Exon1
1
6
Chr18
1,901 bps
1
6
Pard6g
1
6
Chr18
Cre+
Cre+ Up1 405 bps
Cre+ Up3 376 bps
Cre-
Cre+
32604
C57bl6
H2O
32530
32531
32601
32681
32683
32686
3,000bp -
32604
C57bl6
H2O
3,000bp -
32530
32531
32601
32681
32683
32686
1,500bp -
1,500bp -
850bp -
850bp -
400bp -
400bp -
300bp -
300bp -
100bp -
100bp -
WT= 1,833 bps (not detactable)
Flox= 1,901 bps (not detactable)
Cre+ Up1= 405bp
Cre+ Up3 = 376bp
